# Multi-omics analyses identify transcription factor interplay in corneal epithelial fate determination and disease

**DOI:** 10.1101/2022.07.13.499857

**Authors:** Jos GA Smits, Dulce Lima Cunha, Maryam Amini, Marina Bertolin, Camille Laberthonnière, Jieqiong Qu, Nicholas Owen, Lorenz Latta, Berthold Seitz, Lauriane N Roux, Tanja Stachon, Stefano Ferrari, Mariya Moosajee, Daniel Aberdam, Nora Szentmary, Simon J. van Heeringen, Huiqing Zhou

**Author notes:** Corresponding author (HZ).

## Abstract

The transparent corneal epithelium in the eye is maintained through the homeostasis regulated by limbal stem cells, while the non-transparent epidermis relies on epidermal keratinocytes for renewal. Despite their cellular similarities, the precise cell fates of these two types of epithelial stem cells, which give rise to functionally distinct epithelia, remain unknown. We performed a multi-omics analysis of human limbal stem cells from the cornea and keratinocytes from the epidermis, and characterized their molecular signatures, highlighting their similarities and differences. Through gene regulatory network analyses, we identified shared and cell type-specific transcription factors that define specific cell fates, and established their regulatory hierarchy. Single-cell RNA-seq analyses of the cornea and the epidermis confirmed these shared and cell type-specific transcription factors. Notably, the shared and limbal stem cell-specific transcription factors can cooperatively target genes associated with corneal opacity. Importantly, we discovered that *FOSL2*, a direct PAX6 target gene, is a novel candidate associated with corneal opacity, and it regulates genes implicated in corneal diseases. By characterizing molecular signatures, our study unveils the regulatory circuitry governing the limbal stem cell fate and its association with corneal opacity.

## Introduction

Cell fate determination is a complex process essential for normal development and homeostasis. The key role of transcription factors (TFs) in this process has been demonstrated by a plethora of seminal studies where cell conversions can be achieved by forced expression of specific sets of TFs, e.g. generation of induced pluripotent stem cells (1,2). TFs control cell fate determination by regulating the transcriptional program, through binding to cis-regulatory elements (CREs) on the DNA, and by modifying the chromatin environment (3,4). This precise control is essential for tissue integrity and tissue-specific function, and deregulation often leads to pathological conditions (5,6).

The corneal epithelium in the eye and the skin epidermis are two types of stratified epithelia, both derived from the surface ectoderm during embryonic development. The human corneal epithelium is the outermost layer of the cornea, supported by underlying stroma and endothelium, protecting the eye from the outside environment (7–9). It is avascular and transparent, which allows the light into the eye. The proper structure and function of the corneal epithelium are maintained by limbal stem cells (LSCs), located in the limbus, at the rim of the cornea. Differentiating LSCs move centrally to form basal epithelial cells, and stratify to form differentiated epithelial layers (10). Similar to the corneal epithelium in barrier function, the skin epidermis, on the other hand, is non-transparent. The homeostasis of the epidermis is controlled by keratinocytes (KCs) in the basal layer of the epidermis. Basal KCs differentiate vertically and outwards to form different strata of the epidermis (7). Both LSCs and KCs are similar in their cellular morphology, even indistinguishable when cultured *in vitro,* and share expression of basal epithelial genes such as *KRT5* and *KRT14*. Nevertheless, their cell fates are intrinsically distinct, as they initiate and maintain specific epithelial differentiation programs that give rise to the transparent corneal epithelium and non-transparent epidermis, respectively. Insights into the comparison between cell fates of LSCs and KCs will shed light on the control mechanism of their cellular function and related pathological conditions, e.g., corneal opacity. So far, however, the cell fate similarities and differences between KCs and LSCs controlled by TFs and their associated epigenetic mechanisms are not yet understood.

In KCs and the epidermis, key TFs have been studied extensively, both *in vitro* and *in vivo* (11–14). Key TFs include p63, GRHL family proteins, KLF4 and ZNF750, which all regulate transcriptional programs important for KC proliferation and differentiation (12,13,15,16). Many of these TFs, often working together, are known to modulate the chromatin landscape through enhancers (12,13,16). The TF p63 encoded by *TP63* is a key regulator of stratified epithelia and is important for the commitment, proliferation, and differentiation of KCs (13). It binds mainly to enhancers and maintains the epigenetic landscape for the proper epidermal cell identity (16–18). Mutations in *TP63* are associated with developmental disorders like ectrodactyly, ectodermal dysplasia, and cleft lip/palate (EEC) syndrome (OMIM 604292). In EEC, patients present defects in ectodermal derivatives, e.g., epidermis, hair follicles, and nails, but also in other epithelium-lined tissues such as the cornea (19–21). The disease phenotypes of *TP63* mutation-associated disorders are consistent with p63 expression in stratified epithelia (19–21). It has been shown that loss of the typical epidermal identity due to rewired epigenetic circuitry is characteristic of KCs carrying *TP63* mutations associated with EEC (16).

As compared to the wealth of molecular insights of TFs in KCs, the control mechanism of TFs in the corneal epithelium and LSCs is less understood. One of the better-studied TFs is the eye master regulator PAX6. PAX6 is essential for the specification and determination of different parts of the eye, including the retina, iris, lens, and cornea (22–24). In the retina and lens, PAX6 interacts with chromatin modifiers such as EZH2, and cooperates with and regulates other TFs to define cell fates (25–27). In LSCs of the cornea, PAX6 binds to enhancers, together with TFs such as RUNX1 and SMAD3, important for controlling the LSC identity (23,28–32).

Mutations and deregulation of *PAX6* are associated with aniridia (OMIM 106210), a disorder initially characterized by an absent or underdeveloped iris, among other phenotypes such as defects in the retina, pancreas, and neurological systems (33), which is consistent with PAX6 expression in these tissues and organs (24). Relevant to the cornea, up to 90% of aniridia patients show progressive limbal stem cell deficiency (LSCD) leading to corneal opacities (31,34). Interestingly, LSCD and corneal opacities are also present in over 60% of patients with *TP63* mutation-associated EEC syndrome (35,36). In addition to PAX6 and p63, another TF that has been associated with corneal abnormalities is FOXC1, of which mutations are involved in the spectrum of anterior segment dysgenesis, including Peters anomaly and Axenfeld-Rieger syndrome (OMIM 602482) (37). FOXC1 is expressed in the epithelium, stromal, and endothelial cells of the cornea, and is shown to be upstream and regulating PAX6 (38,39). Recently, two reports suggested that loss of *PAX6* or *FOXC1* in LSCs gives rise to loss of the LSC identity, and these PAX6 or FOXC1 deficient LSCs acquire a KC-like cell signature, indicated by upregulated expression of epidermal stratification marker genes (30,39,40). How TFs like PAX6, p63, and FOXC1 regulate their target genes in LSCs and how their mutations give rise to LSCD and corneal opacities are not yet fully understood. Therefore, a comprehensive characterization and comparison of molecular signatures between LSCs and KCs will not only identify shared and tissue-specific TFs controlling cell fates but also provide insights into the pathomechanisms of LSCD and other corneal opacity disease mechanisms.

In this study, we performed in-depth analyses of the transcriptome and the epigenome of human LSCs and KCs cultured *in vitro*, and characterized differentially expressed genes and regulatory regions between the two cell types. Using a gene regulatory network-based method, we identified key TFs and their hierarchy controlling epithelial programs that are shared by KCs and LSCs, and those that are distinct for each cell type. Expression patterns of the key TFs were further validated with *in vivo* single-cell RNA-seq data from the cornea and the epidermis. Importantly, we showed that the key TFs and their target genes that drive the LSC specific epithelial program are associated with corneal diseases, and identified a novel disease candidate *FOSL2* associated with corneal opacity.

## Results

### Distinct epithelial gene expression patterns define cell fate differences of skin keratinocytes and cornea limbal stem cells

To characterize gene expression patterns that define the cell fate difference between human cornea limbal stem cells (LSCs) and human skin keratinocytes (KCs) (Fig 1A), we used cultured LSCs established from post mortem limbal biopsies and basal KCs from skin donors. Both cultured cells have the capacity to re-generate stratified epithelial tissues *in vitro* (41,42), and have high p63 expression (SFig 1F), thus exhibiting the progenitor cell state. We performed comparative RNA-seq analyses from bulk and pseudobulk RNA-seq data (aggregated from single-cell RNA-seq (scRNA-seq) using cultured LSCs and KCs) experiments and incorporated our data with publicly available RNA-seq data for robustness (supplementary Table 1) (supplementary Fig 1) (16,32). Single-cell data was aggregated because no measurable heterogeneity was detected in these cultured cells, except for cell cycle differences (supplementary Fig 1E).

**Fig 1.**
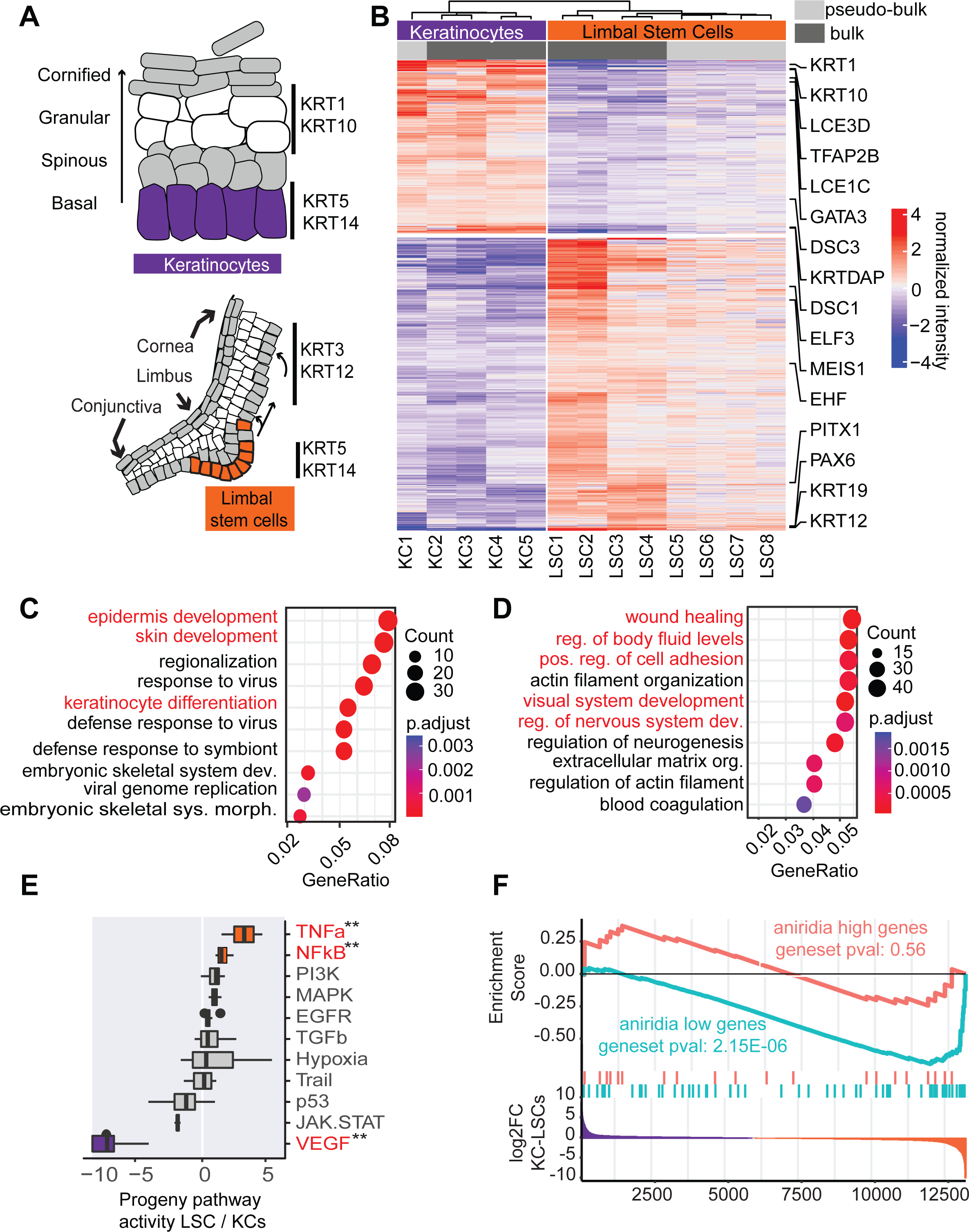
RNA-seq analysis of LSCs and KCs. (A) Schematic picture of the epidermis and the limbus. (B) Heatmap of normalized expression of differentially expressed genes between LSCs and KCs (adjusted pval < 0.01, log2 FC > 1.5). Differentially expressed genes are clustered using k-means clustering with 2 clusters. (C) GO-term enrichment of KC-high genes. (D) GO-term enrichment of LSC-high genes. (E) PROGENy pathway activity analysis, with scores sorted based on the LSC/KC ratio. Pathways depicted in red are differential, the color is grey if non-differential, orange if higher in LSC, and purple if higher in KCs. (F) Gene Set Enrichment Analysis of differentially expressed genes identified in aniridia patient LSCs, as compared to controls, up-and down-regulated genes (aniridia high and low, respectively) were tested for enrichments against the KC-LSC foldchange.

Through pairwise comparison, we identified 1251 differential expressed genes between LSCs and KCs. Among them, 793 genes had higher expression in LSCs (referred to as LSC-high genes), while 459 differential genes were more highly expressed in KCs (referred to as KC-high genes). This analysis resulted in typical genes for both epithelial cell types: LSC-high genes with limbal and corneal epithelial genes including KRT19 and the eye master regulator PAX6 (Fig 1B, supplementary Fig 1F), whereas KC-high genes contained epidermal markers such as *KRT1*, *KRT10*, *LCE3D* & *LCE3C*. Although some of these detected genes are associated with epithelial stratification, e.g., *KRT3* and *KRT12* for the cornea and *KRT1* and *KRT10* for the epidermis, their expression was significantly lower (5-20 fold) than their expression in stratified epithelial cells (16,39) (supplementary Fig 1G), confirming the progenitor states of the cultured LSCs and KCs. It should be noted that TP63 is highly expressed in both LSCs and KCs (supplementary Fig 1F), and therefore is not identified as differential.

Gene Ontology (GO) enrichment analysis (43,44) of KC-high genes identified enrichment of GO terms related to the “epidermis” and “skin development” (Fig 1C). GO terms associated with “response to virus” were enriched due to detected immune and interferon-related genes among KC-high genes, consistent with the gene set enrichment analysis (GSEA) using the hallmark gene set of the MsigDB collection (45,46) that identified enrichment of the interferon-alpha and gamma response (supplementary table 4). Finally, PROGENy pathway target gene analysis (47) identified higher VEGF signaling in KCs (Fig 1E), indicating that VEGF vascularization-related genes are lowly expressed or completely repressed in LSCs. This is in line with the avascularized state of the cornea. LSC-high genes were enriched for terms such as “wound healing” and “positive regulation of body fluid levels” as well as “visual system development” and “regulation of nervous system development” (Fig 1D). PROGENy analysis identified the TNF-α and NFKβ pathways associated with LSCs (Fig 1E), such as *CXCL1,3,5,6* and *TNF-α*. Consistently, TNF-α and NF-kB signaling was also identified by KEGG pathway (48), GO and GSEA analyses using the hallmark gene set of the MsigDB collection (supplementary Fig 2). Finally, GSEA enrichment using the C8 dataset of the MsigDB collection that contains single-cell dataset gene lists also identified enrichment for genes in the “Descartes fetal eye corneal and conjunctival epithelial cells” (supplementary table 4) with more signal in the LSCs.

Next, we asked the question of whether LSCD-associated LSCs acquire a KC-like cell fate. To test this, we examined the cell fate of LSCs from patients with aniridia, a disease mostly caused by *PAX6* haploinsufficiency. We performed RNA-seq analysis of primary LSCs of two aniridia patients and post-mortem cornea-extracted LSC controls. Aniridia and control LSCs were both expanded in KSFM medium conditions and were on passage three when processed. These data were integrated with other aniridia RNA-seq data published previously using the same procedures and culture conditions (49) (supplementary Table 1). We obtained 73 differentially downregulated genes (aniridia low genes) and 22 upregulated genes (aniridia high genes) in aniridia patient LSCs, as compared to LSCs from healthy controls (supplementary Fig 3). Aniridia low genes included the *PAX6* target gene *KRT12* and other corneal and epithelial genes such as *TGFBI*, *CLND1*, *GJB6*, *IL36G*, *LAYN*, *NMU,* and *TMEM47.* Many of these epithelial genes are potential PAX6 target genes reported in an immortalized LSC model where one allele of *PAX6* was deleted (50). We then applied GSEA to compare them to LSC and KC gene expression signatures, to investigate whether these deregulated genes due to *PAX6* haploinsufficiency represent the changed cell fate of these aniridia LSCs. Indeed, as expected, aniridia low genes were enriched among genes expressed highly in LSCs (P-value 2.8E-05) (Fig 1F), indicating a loss of the LSC cell fate in aniridia LSCs. Among aniridia high genes, we found *GATA3* present in genes expressed highly in KCs (supplementary Fig 3). Nevertheless, there was no detected enrichment of aniridia high genes among genes expressed highly in KCs representing the KC cell fate, arguing against the postulated model that PAX6-deficient LSCs acquire a KC-like cell fate at the transcriptome level. These data suggest that additional mechanisms, such as TFs other than PAX6, contribute to the cell fate difference between KCs and LSCs.

### The epigenetic states of cis-regulatory elements correlate with gene expression patterns

To understand the mechanisms underlying different cell fate controls of LSCs and KCs, we identified cis-regulatory elements (CREs) and their epigenetic states that drive gene expression differences. We generated an extensive multi-omics dataset of LSCs and KCs and integrated these with other published data (supplementary Fig 4A, supplementary Table 2, and Table 3).

The complete dataset included ATAC-seq for open chromatin regions representing CREs (32) and ChIP-seq of histone modifications, H3K27ac and H3K4me3 marking active CREs, and H3K27me3 that marks repressed chromatin regions (Fig 2A) (17,32). Using ATAC-seq analysis, we identified 124,062 CREs in the two cell types. Approximately 80% of these CREs were accessible in both cell types (supplementary Fig 5). The public ATAC-seq data was compared to the data generated using Pearson correlation (supplementary Fig 5F). This comparison revealed a distinct biological difference between KCs and LSC, which was more prominent than technical variation across different laboratories and techniques.

**Fig 2.**
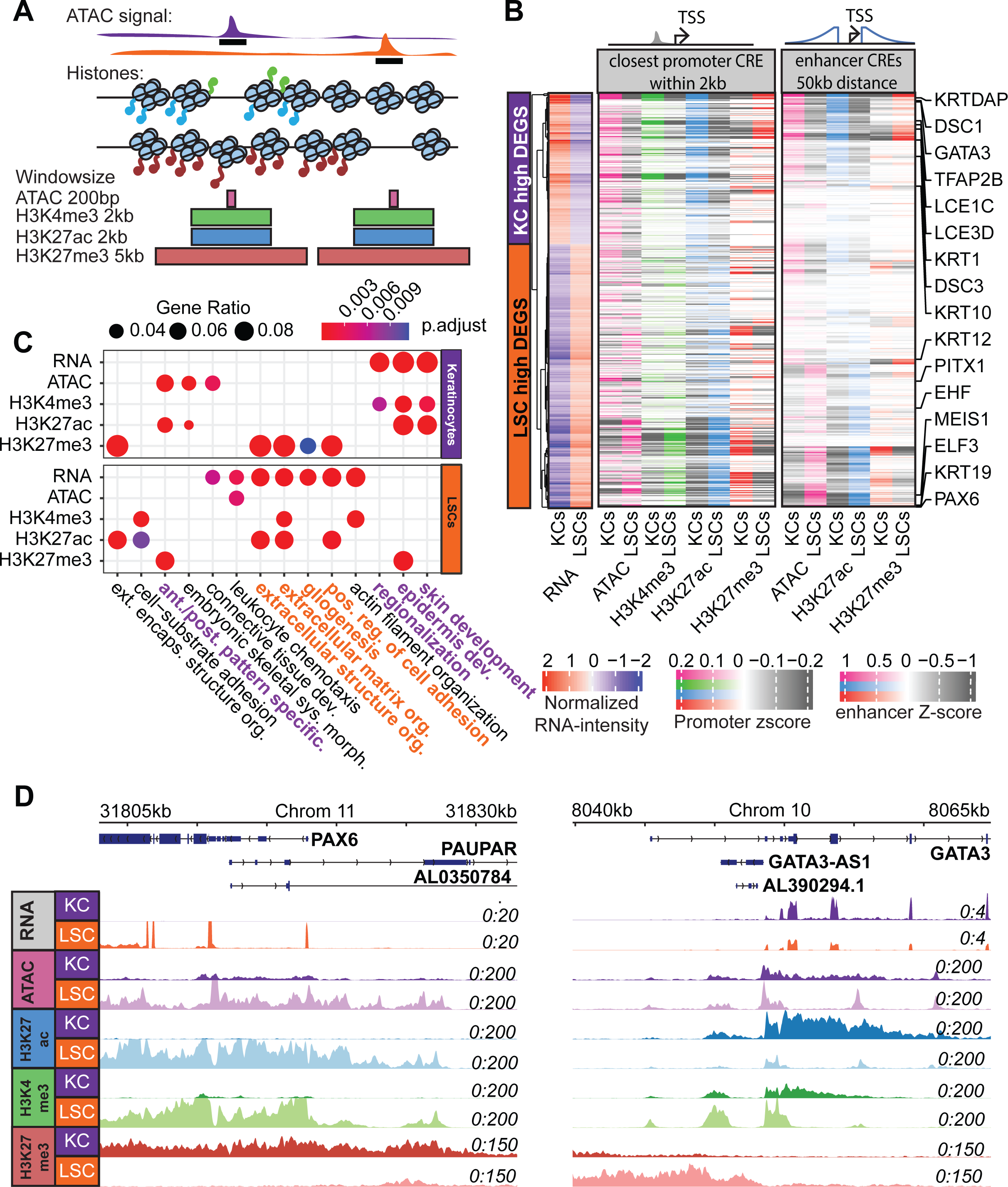
Cis Regulatory Element (CRE) analysis. (A) Schematic overview of CRE identification and quantification. Signals of each analysis were quantified by different window sizes covering the ATAC-seq peak summit. (B) Heatmap of the Z-scores of the quantile normalized ATAC-seq and histone mark signals near LSC- and KC-high genes. For promoter CREs, it corresponds to the closest CRE within 20kb to the transcription start site (TSS). For enhancer CREs, the signals of all CREs within a 100kb window near a TSS were quantified, distance weighted, and summed (C) GO-term enrichment of LSC- and KC-high genes and genes close (within 20kb) to differential CREs. (D) PAX6 and GATA4 TSS loci with signals of RNA-seq, ATAC-seq, ChIP-seq of H3K27ac, H3K4me3, and H3K27me3 in KCs and LSCs

To examine the differential epigenetic states of these CREs in LSCs and KCs, we quantified ATAC-seq and histone modification signals in windows covering these CREs (Fig 2A). This resulted in 35,348 CREs with differential epigenetic signals, about 28.5% of CREs (supplementary Fig 5). To assess the correlation between these differential CREs and the expression of their nearby genes, we considered both CREs at the promoter regions (promoter CREs) and enhancer CREs located within 50kb-distance from the genes (enhancer CREs) (Fig 2B, supplementary Fig 4B). As expected, high ATAC, H3K4me3, and H3K27ac signals correlated with high gene expression, while the strong signals of repressive H3K27me3 correlated well with lowly expressed genes or genes with undetectable expression in the corresponding cell types, e.g., the loci of *PAX6*, *GATA3*, *HOXA9* and *TNF-α* (Fig 2B, 2D, supplementary Fig 4D). The strong repression signals marked by H3K27me3 in KCs at genes that are key for the LSC fate such as *PAX6* suggest a repression mechanism in KCs to prevent inappropriate gene expression that defines the LSC fate. All annotated CRE regions are available on a publicly accessible trackhub (see Data availability).

Next, we performed GO analysis on genes that are close to differential CREs (Fig 2C). For H3K27ac and H3K4me3 that mark active CREs, GO terms such as “epidermis” and “skin development” were identified for CREs with strong signals in KCs. Furthermore, “positive regulation of cell adhesion” and “extracellular matrix organization” terms were found for CREs with strong H3K27ac and H3K4me3 signals in LSCs. This is consistent with identified differentially expressed genes in each corresponding cell type. In contrast, the repressive mark H3K27me3 is anti-correlated with gene expression (Fig 2C).

Furthermore, in line with the enrichment of *TNF-α* and *NF-kB* signaling pathways in LSCs identified by PROGENy analysis of differentially expressed genes, higher H3K27ac, and H3K4me3 signals were present in the loci of *TNF-α* and *NF-kB* target genes in LSCs, as compared to in KCs, while these loci in KCs were repressed by H3K27me3 (supplementary Fig 4C).

### Gene regulatory network analysis identifies transcription factors controlling distinct epithelial cell fates and their hierarchy

Using the identified differential CREs, we set out to identify key TFs driving the cell fate differences between LSCs and KCs. TF binding motif enrichment was performed using Gimme Motifs (51) in all differential CREs marked by ATAC, H3K4me3, H3K27ac, and/or H3K27me3 signals. In general, TF motifs enriched in CREs with active marks, ATAC, H3K4me3, or H3K27ac, in one cell type, were also enriched within regions with the repressive H3K27me3 mark in the other cell type (Fig 3A, supplementary Fig 6A). For example, TF motifs that are linked to FOXC1, TEAD1, JUN, PAX6, FOS, RUNX2, OTX1, ELF3, SOX9, and REL were detected in differential CREs marked by high active mark signals in LSCs but also marked by high H3K27me3 in KCs. Consistent with our expectation, the enrichment of the PAX6 motif in differential CREs with higher active mark signals was detected in LSCs, as PAX6 is specific for LSCs but not for KCs. As REL is a TF involved in TNF-α and NF-kB pathways, the detection of the REL motif is consistent with the enrichment of TNF-α and NFKβ signaling genes among LSC-high genes. Notably, the FOS motif that is associated with FOS, FOSL1, FOSL2, and JUN (supplementary Fig 6A) was present in approximately 10% of all variable CREs in LSCs, the highest among all motifs (Fig 3A). Motifs enriched in KC active CREs included those linked to KLF6, GRHL1, HOXC10, GATA3, NFIA, CEBPA, and CTCF. These motifs were also enriched in CREs marked by high H3K27me3 signals in LSCs. Enriched motifs could mostly be linked to TFs with high expression differences between LSCs and KCs, e.g., FOXC1, PAX6, and FOS are highly expressed in LSCs, while HOXC10, GATA3, and CEBPA are highly expressed in KCs (Fig 3A).

**Fig 3.**
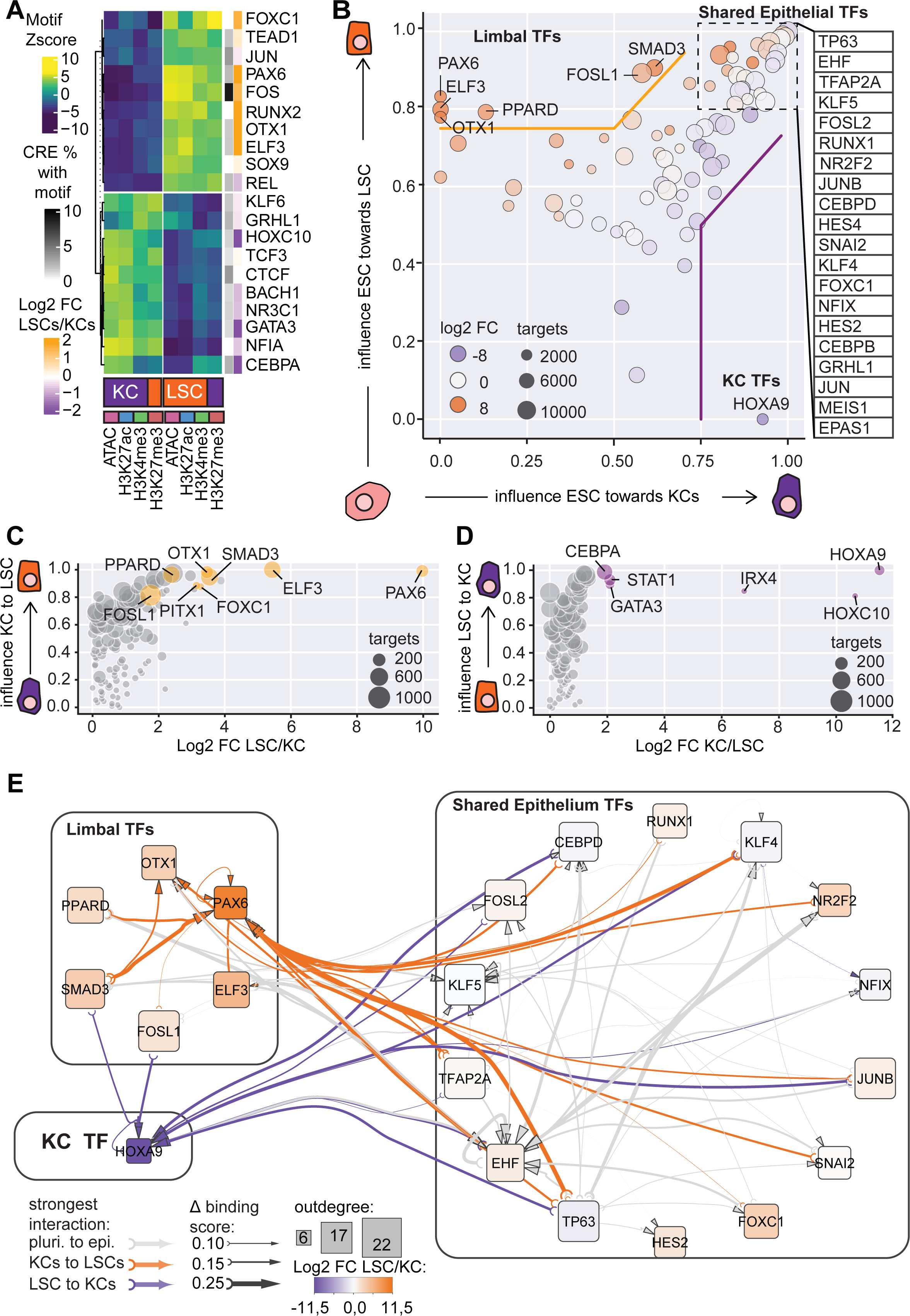
TFs and TF hierarchy controlling distinct epithelial cell identity. (A) Heatmap of motif enrichment Z-scores detected in variable CREs and the corresponding TFs. The percentage of CREs containing the motifs and the expression ratio of TFs in LSCs and KCs are indicated. (B) ANANSE influence score plot of TFs identified in ESC-KC (x-axis) and ESC-LSC (y-axis) comparison. Circle size represents the maximum number of target genes of a TF. The color represents log2FC between LSC/KC (orange LSC high; purple KC high). (C) ANANSE influence score plot of TFs identified in KC-LSC comparison. (D) ANANSE influence score plot of TFs identified in LSC-KC comparison. (E) TF hierarchy is indicated by the binding score of a TF to its target TF locus, and the cell type specific regulation is indicated by the binding score difference of the TF at the target TF locus between cell types. When a binding score difference in KC-LSC comparison is greater than the mean of the difference in ESC-KC and ESC-LSC comparison, this TF regulation of the target TF is annotated as either KC-(purple arrows) or LSC specific (orange arrows) regulation. Otherwise, the regulation is annotated as ‘shared regulation’ for both cell types (grey arrows). The degree of binding score difference is indicated by the thickness of the arrows. Outdegree node size represents the number of target genes. Fold change of TF gene expression in LSC and KCs is represented by orange (LSC-high) and purple (KC-high) colors.

To predict gene regulation by considering the expression of both TFs and their targets, we applied ANANSE (52), a gene regulatory network method to identify key TFs for cell identities. ANANSE integrates CRE activities and TF motif predictions with the expression of TFs and their target genes, to generate a gene regulatory network of the specific cell type. Subsequently, a pairwise comparison of gene regulatory networks from two cell types is performed to identify the most influential TFs that differentiate between the two cell types. The overall importance of identified TFs is represented by their influence score, scaled from 0-1 (Material and Method) (52).

Because many key TFs for LSC and KC fates that are shared in these two cell types such as p63 had similar gene expression levels (supplementary Fig 1F), these TFs could not be identified through a direct comparison of the differential gene regulatory network between these two cell types using ANANSE. Therefore, we decided to utilize embryonic stem cells (ESCs), a completely different cell type as compared to either LSCs or KCs, as the reference point for this differential network analysis. This enabled us to identify not only distinct but also shared TFs for LSC and KC fates. Using RNA-seq, ATAC-seq, and H3K27ac data from ESCs (53), we performed a pairwise comparison of gene regulatory networks between ESCs versus KCs or ESCs versus LSCs. When predicting the TFs driving the LSC or KC fates from ESCs, ANANSE resulted in 70 epithelial TFs that had influence scores above 0.5 in both ESC-LSC and ESC-KC pairwise differential network analysis. Many TFs are known to be important for epithelial cell function, such as TP63, EHF, TFAP2A, TFAP2C, FOSL2, the KLF family protein 3,4,5,6,7, JUNB, CEBPD, CEBPB, and RUNX1. We classified these TFs as shared epithelial TFs (Fig 3B, supplementary Fig 7). Intriguingly, when conducting a network outdegree analysis, that quantifies the number of targets of a TF within the top 20 TFs, FOSL2, JUN, TP63, and TFAP2A were identified as regulating most other TFs. This implies their importance in driving other TF expression (supplementary Fig 7B, C). The highest outdegree of FOSL2 is in line with the high percentage of detected FOS motif (Fig 3A).

Importantly, in the ESC-LSC and ESC-KC differential network analysis, TFs with high influence scores in LSCs but with undetectable (PAX6, ELF3, OTX1, PPARD) or low (FOSL1 and SMAD3) influence scores in KCs were considered as LSC specific TFs (Fig 3B). Direct pairwise differential network analysis between LSCs and KCs (Fig 3C) was largely consistent with these findings. Interestingly, FOXC1 was annotated as a shared epithelial TF, whereas in the direct KC-LSC comparison, it was identified as a specific TF for the LSC fate. This is probably due to the higher expression of FOXC1 in LSCs. For KC specific TFs, we only identified HOXA9 in the ESC-LSC and ESC-KC differential network analysis (Fig 3B). The direct pairwise differential network analysis between KCs and LSCs (Fig 3D) confirmed this. Next to HOXA9, the direct comparison identified other TFs such as GATA3, IRX4, and CEBPA with high influence scores in KCs, indicating that KC-LSC pairwise comparison is more sensitive for detecting KC specific TFs.

Finally, we set out to dissect the TF regulatory hierarchy for the cell identity differences between LSCs and KCs, by identifying potential target TFs of the shared and specific TFs. For this analysis, we did not consider the expression level of TFs themselves, to ensure that the prediction is mainly driven by the potential binding of a TF to its target loci, represented by the binding score. If a target TF is regulated by a TF with similar binding scores in both LSCs and KCs compared to ESCs, this regulation is annotated as a ‘shared regulation’; if the binding score is significantly higher in one cell type than in the other (Material and Method), the regulation of the TF-target TF pair is annotated as ‘cell type specific regulation’. We included the top shared and specific TFs, 15 shared epithelial TFs, and six LSC specific TFs (PAX6, ELF3, OTX1, PPARD, SMAD3, and FOSL1), and the KC specific TF HOXA9 (Fig 3B, 3E, supplementary Fig 6B, 6C, 6D). As expected, many shared TFs are regulating each other via ‘shared regulation’ (Fig 3E, grey arrows). Consistently, cell type specific TFs regulate their target TFs via ‘cell type specific regulation’ (Fig 3E, orange arrows), e.g., *PAX6* is predicted to be regulated by SMAD3 and PPARD. Furthermore, many autoregulation loops were also detected, e.g., PAX6 in LSCs and HOXA9 in KCs. We also found that shared TFs may regulate cell type specific TFs in via ‘cell type specific regulation’. For example, p63, FOXC1, and TFAP2A were identified as shared TFs between KCs and LSCs, but they were predicted to regulate *PAX6* in LSCs. In summary, our molecular characterization using KCs and LSCs cultured *in vitro* identified shared and cell type specific TFs for the LSC and KC fates. p63, FOSL2, EHF, TFAP2A, KLF5, RUNX1, CEBPD, and FOXC1 are among the shared epithelial TFs for both LSCs and KCs. PAX6, SMAD3, OTX1, ELF3, and PPARD are LSC specific TFs for the LSC fate, and HOXA9, IRX4, CEBPA, and GATA3 were identified as KC specific TFs. Furthermore, LSC and KC fates are defined by cooperative regulation of both shared and cell type specific TFs.

### Single-cell RNA-seq analysis of the cornea and the epidermis validates the expression of key transcription factors controlling cell fates

Since our multi-omics analysis was performed on LSCs and KCs cultured *in vitro,* we assessed single-cell RNA-seq datasets derived from the cornea and the epidermis, to confirm that the molecular signatures of LSCs and KCs in our study indeed represent those of epithelial stem cells maintaining the corneal limbus and the epidermis (54,55). By clustering single cells according to marker gene expression (supplementary Fig 8), we selected the cell clusters corresponding to the stem cells as pseudobulk for further differential gene expression analysis. For the epidermis, we selected cells with high *KRT14*, *KRT5,* and low *KRT1* and *KRT10* expression as basal KCs, and for the cornea, cells with high *S100A2*, *PAX6,* and *TP63* expression and without *CPVL* expression as LSCs, because *CPVL* has been proposed as a marker with neural crest origin (55).

Consistent with the *in vitro* findings, the *in vivo* LSCs expressed high levels of *PAX6*, *ELF3*, *FOXC1*, *FOSL1, OTX1,* and *SMAD3*, whereas the *in vivo* KCs expressed high levels of *HOXA9, CEBPA, and GATA3* (Fig 4A). GO analysis identified similar functions of differentially expressed genes between the *in vivo* LSCs and KCs, as compared to those from *in vitro* cultured cells (supplementary Fig 8E, F). Furthermore, PROGENy analysis of differentially expressed genes between *in vivo* LSCs and KCs showed that TNF-α and NF-kB pathway genes are significantly enriched in *in vivo* LSCs, e.g., *CXCL1,2,3,8,20* and *NFKB1* (Fig 4B). GSEA analysis using the hallmark gene set of the MsigDB collection also identified enrichment for TNF-α signaling genes (supplementary table 5).

**Fig 4.**
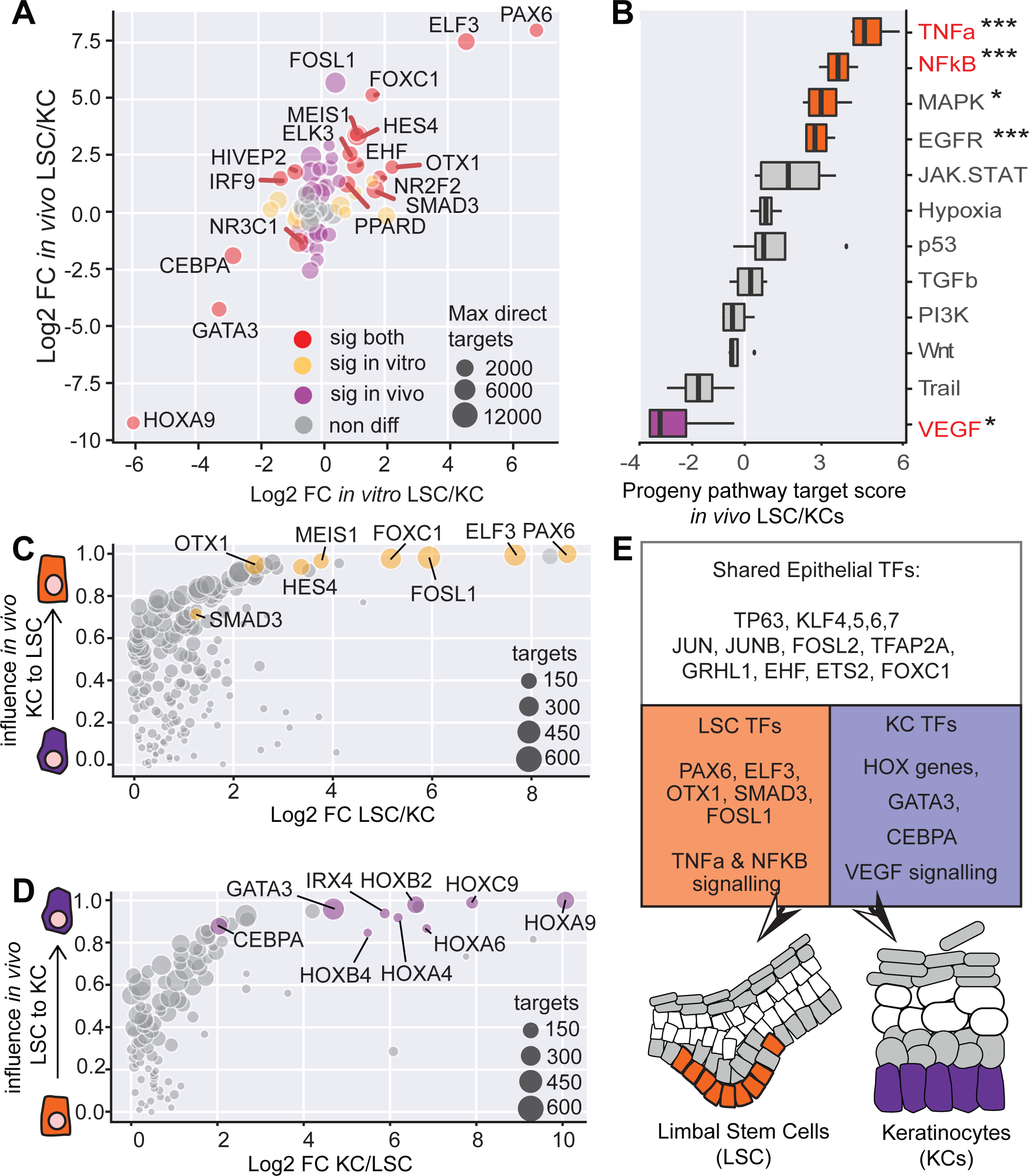
Validation of key TF expression using in vivo single-cell RNA-seq. (A) Fold change comparison of identified TFs using in vivo and in vitro data. (B) PROGENy pathway analysis of in vivo LSCs and KCs. (C) ANANSE influence score plot of in vivo basal KCs to LSCs. (D) ANANSE influence score plot of in vivo LSC to basal KCs. E) Summary of the identified shared and specific TFs.

As the data of CREs of *in vivo* tissues were not available, we utilized the *in vitro* ATAC and H3K27ac datasets for differential gene regulatory network analysis, together with the *in vivo* pseudobulk data used for gene expression analysis above. Here we assumed that since the GRN analysis is largely driven by gene expression data, this analysis assessed the influence of TFs on *in vivo* LSC and KC fate differences. Overall, the *in vivo* data identified similar cell type specific TFs, as compared to *in vitro* cultured cells (Fig 4C and 4D). In *in vivo* LSCs PPARD however was not detected, PAX6, ELF3 FOXC1, and FOSL1 exhibited the highest influence scores, and TFs with the highest influence scores in *in vivo* basal KCs were HOX TFs and a few others such as CEBPA, GATA3, and IRX4.

Taken together, our analyses showed clear consistency between *in vivo* and *in vitro* derived data and identified key TFs driving the cell fate of between LSCs and KCs (Fig 4E).

### FOSL2 is a novel transcription factor controlling the LSC fate and associated with corneal opacity

As several key TFs defining the LSC fate, either shared or cell type specific, such as p63 and PAX6, respectively, are associated with corneal opacity, we explored the potential involvement of LSC TFs in the pathomechanism of corneal diseases. For this we leveraged the whole genome sequencing data in the 100,000 Genomes Project at Genomics England UK to identify variants of uncertain significance that may have functional consequences. For establishing a suitable cohort, we identified a total number of 33 unsolved participants with human phenotype ontology (HPO) terms associated with corneal opacity (supplementary table 8). Next, we screened for variants in the coding regions of the top 15 shared epithelial TFs as well as the six LSC specific TFs identified in our study. In a proband with band keratopathy (HP:0000585), we identified a rare *de novo* heterozygous missense variant in *FOSL2* (2:28412095:C:T, genome build GRCh38/hg38, NM_005253.4:c.628C>T) (genomAD allele frequency 0.0000922) (supplementary Fig 9A), giving rise to a predicted damaging amino acid change (NP_005244.1:p.(Arg210Cys)), based on most major prediction tools (supplementary Fig 9A, supplementary table 9).

As the role of FOSL2 in LSCs is completely unknown, we first investigated the protein expression of FOSL2 within the cornea. The immunostaining results showed that FOSL2 protein is expressed in both limbus and central cornea (Fig 5A, supplementary Fig 9B), co-localizing in the nucleus with other LSC TFs, e.g. p63, PAX6 and its closely related AP1 complex factor FOSL1 (supplementary Fig 9C-F). The same was also visible in LSCs (and KCs, with the exception of PAX6) cultured *in vitro* (supplementary Fig 9G-J). The identified variant associated with corneal opacity, together with the expression pattern, indicated an important function of FOSL2 in the cornea.

**Fig 5.**
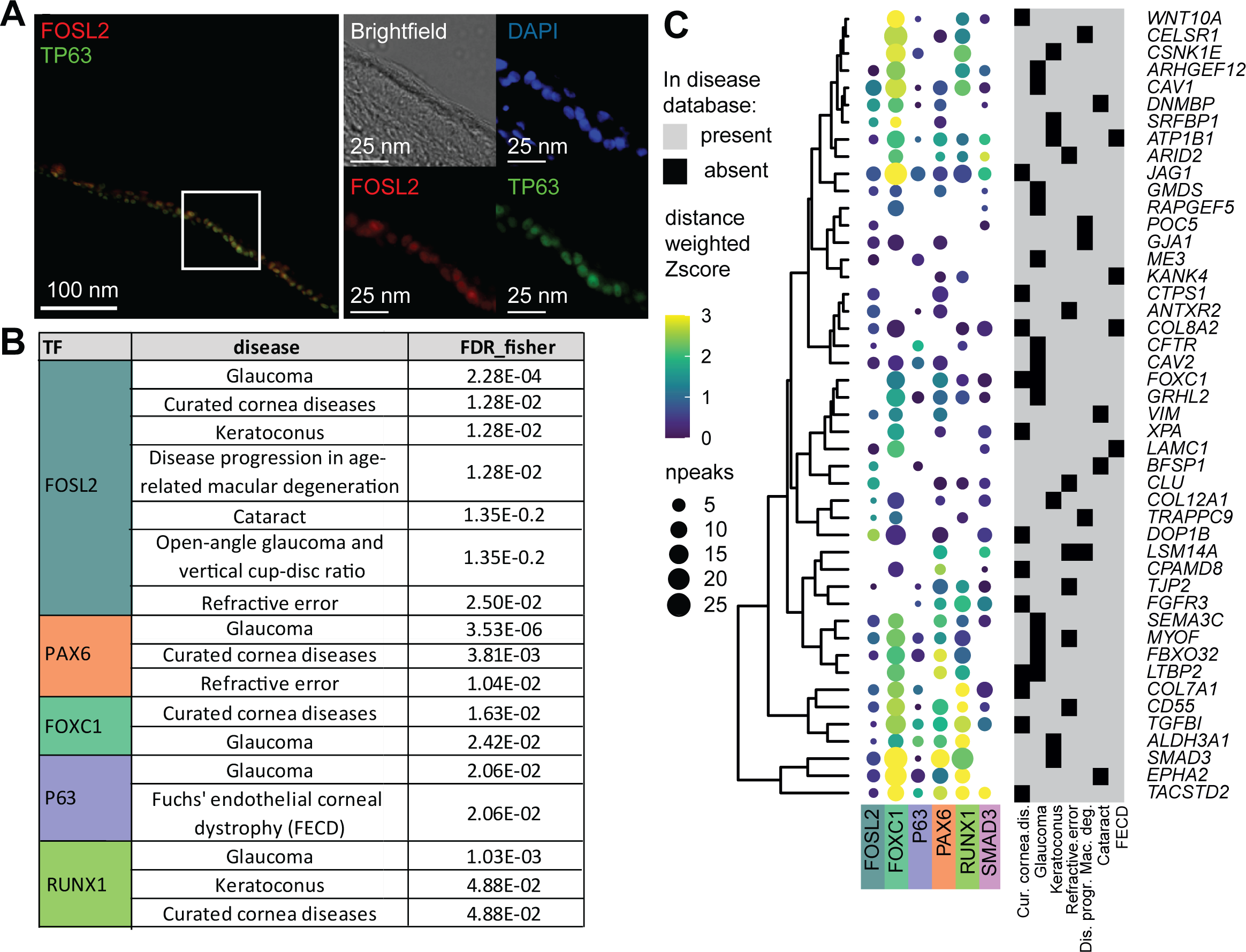
Transcription factors regulation of corneal disease-associated genes. (A) FOSL2 and p63 staining of the peripheral cornea. (B) TFs that bind to gene loci associated with corneal abnormalities with significantly higher occurrence (FDR), as compared to binding to all genes in the whole genome. FDR was calculated with Fisher exact testing. (C) Dot plot showing the TF binding intensity (color bar) and the number of binding peaks (npeaks, dot size) near the disease genes that contain a significant number of potential TF binding. The number of peaks is within a 100kb region of the transcription starting site (TSS); TF binding intensity score is the weighted z score of the quantile log normalized intensities distance weighted. FECD, Fuchs Endothelial Corneal Dystrophy.

To further explore the association of FOSL2 with corneal diseases, we questioned whether the target genes of FOSL2 are enriched for corneal disease genes, similar to other known LSC TFs such as PAX6, FOXC1, and p63. The rationale behind this question is based on the concept that many disease-associated TFs could contribute to the diseases not only because of their mutations but also due to abnormal regulation of their target genes. For this analysis, we employed two types of data, DNA binding profiles of key LSC TFs to predict their target genes and a comprehensive list of corneal disease genes. For TF DNA binding profiles, we generated Cleavage Under Targets and Release Using Nuclease (CUT&RUN) of FOSL2 and ChIP-seq of the p63 protein in LSCs, and additionally incorporated publicly available ChIP-seq data of PAX6, FOXC1, RUNX1 and SMAD3 in LSCs (30,32). Except for FOSL2, all other TFs are known to regulate the LSC fate. To collect genes that are associated with corneal phenotypes, we used two approaches. First, we performed a literature search and constructed a gene list of 161 genes associated with LSCD and inherited corneal opacification diseases, which included epithelial but also stromal and endothelial cornea diseases (e.g., corneal dystrophies, keratoconus) and ocular as well as systemic syndromes with known corneal manifestations (supplementary table 7). In addition to this curated disease gene list, we used disease genes assembled in the EyeDiseases database (56) that includes genes associated with other eye diseases such as “glaucoma” and “refractive error”. To assess whether TF binding of the key TFs to corneal disease gene loci is more probable than random, we mapped TF binding sites to the nearest genes in order to detect a statistically significant enrichment of disease genes bound by TFs (supplementary Fig 10A). With this method, we identified that the genes associated “curated corneal disease” and “refractive error” were significantly more likely to be bound by FOSL2 and PAX6 (Fig 5B and supplementary Fig 10C). Similarly, the genes related to “glaucoma” were enriched in binding by FOSL2, p63, PAX6, FOXC1, and RUNX1 (Fig 5B and supplementary Fig 10C), and genes related to “keratoconus” were enriched in FOSL2 and RUNX1 binding. Additionally, we observed enrichment of FOSL2 binding in genes related to “disease progression in age-related macular degeneration”, “cataract”, “open-angle glaucoma and vertical cup-disc ratio”, FOXC1 binding in “curated corneal disease” and “glaucoma”, p63 binding in “Fuchs’ endothelial corneal dystrophy” (FECD), and RUNX1 binding in genes associated with “keratoconus” and the “curated cornea disease” (supplementary Fig 10C).

The binding signal of these TFs within the TSS regions of the top cornea disease-related genes is summarized by the quantification of binding signals (Fig 5C). In parallel, we investigated the TF-disease association by combining TF binding signals with the distance between the TF binding peaks and the disease genes, and employed a Mann-Whitney U statistical test to identify a potential binding increase (supplementary Fig 10B).This gave rise to similar TF-disease association enrichment results (supplementary Fig 10C)

Overall, our analysis highlighted that FOSL2 and several other key LSC TFs binding is enriched in the TSS regions of cornea disease genes, suggesting that they directly regulate cornea disease genes.

### FOSL2 is a direct PAX6 target gene and regulates angiogenesis and tight junction genes in LSCs

To further assess the function of FOSL2 in LSCs and corneal opacity, we performed siRNA knockdown of *FOSL2* in primary LSCs. Knockdown of *FOSL2* was validated using qPCR and RNA-seq and displayed an efficiency of over 80%, as compared to control siCTR (0.12 fold ±0.068, n=4) (Fig 6A, supplementary Fig 11A,B), and gave rise to 212 differentially expressed genes. Because PAX6 is an undisputed regulator in LSCs and defects are associated with LSCD and corneal opacity, we also performed *PAX6* knockdown (siPAX6). Intriguingly, *FOSL2* expression was significantly downregulated upon *PAX6* knockdown, as compared to siCTR (0.5 fold) (Fig 6A). Furthermore, several high-confidence PAX6 binding peaks were identified surrounding the *FOSL2* locus (Fig 6B), demonstrating that *FOSL2* is a direct target gene of PAX6 in LSCs.

**Fig 6.**
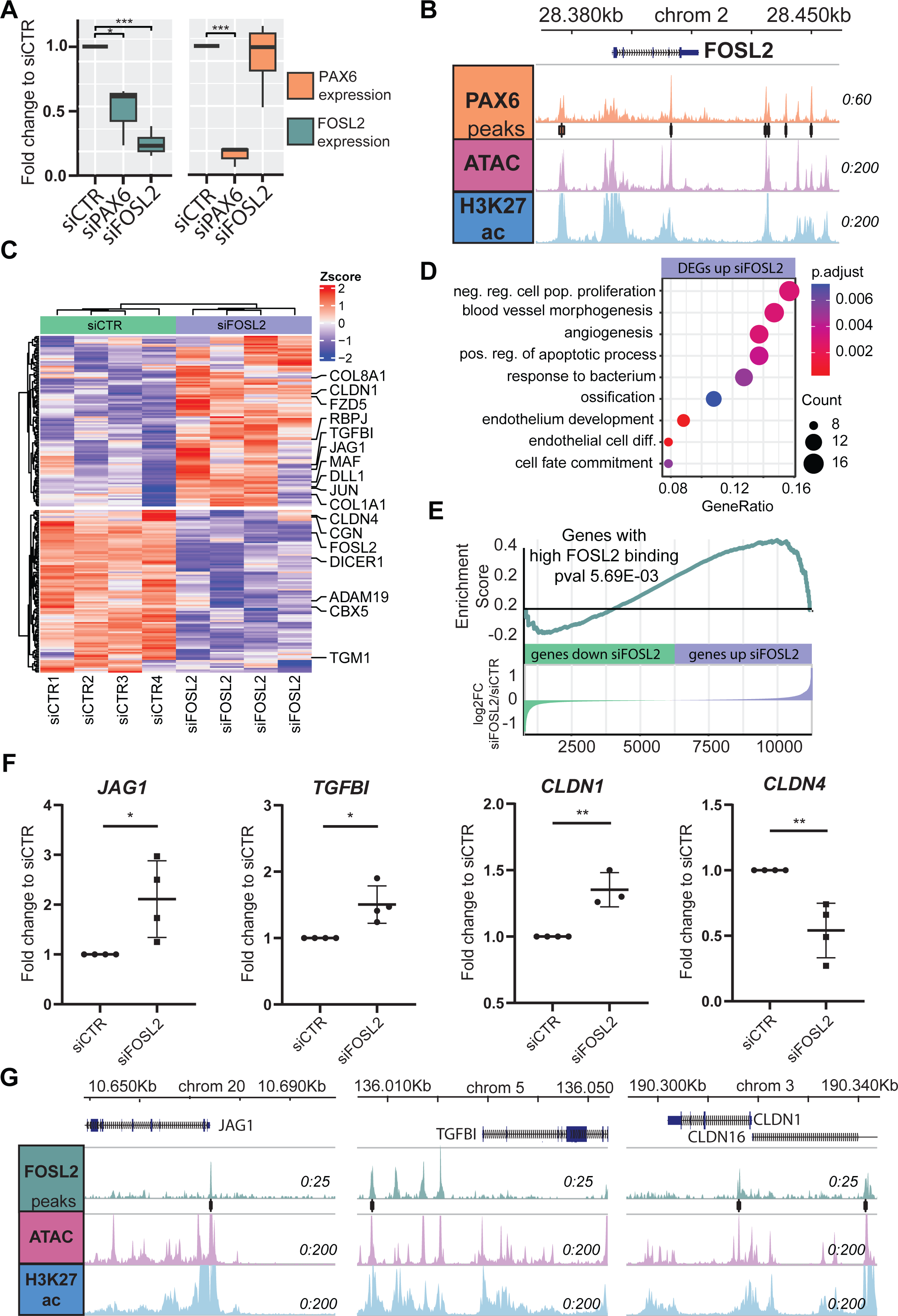
FOSL2 siRNA knockdown results in deregulation of angiogenesis and tight junction genes. (A) Normalized Transcriptome difference of FOSL2 and PAX6 upon knockdown of FOSL2 (siFOSL2) and PAX6 (siPAX6). (* pval<0.05, ** pval<0.01, *** pval<0.001, paired DESEQ2 differential expression testing). (B) FOSL2 TSS locus with PAX6 binding signal, ATAC seq, and H3K27ac ChIP-seq in LSCs. (C) Heatmap of differentially expressed genes in siFOSL2, with Zscore plotted. (D) GO-term enrichment of genes upregulated in siFOSL2 (E) GSEA enrichment analysis based on the expression foldchange (siFOSL2/siCTR) of genes with nearby FOSL2 binding signal. (F) Expression of JAG1, TGFBI, CLDN1, and CLDN4 were measured in control LSCs (siCTL) and FOSL2 siRNA-knock down (siFOSL2) samples (n=4). Values represent the fold change difference of siFOSL2/siCTR and were normalized to internal housekeepers GAPDH and ACTB. (* pval<0.05, ** pval<0.01, *** pval<0.001, unpaired t-test analysis). (G) JAG1, TGFBI, and CLDN1 TSS loci with FOSL2 CUT&RUN signal, ATAC-seq, and H3K27ac ChIP-seq in LSCs.

Among the 212 differentially regulated genes detected in siFOSL2 (Fig 6C), 108 were upregulated, with significant enrichment of genes involved in “negative regulation of cell population proliferation”, “blood vessel morphogenesis”, “angiogenesis”, and “endothelial cell differentiation” (Fig 6D). Furthermore, predicted FOSL2 target genes based on the nearby FOSL2 binding sites were enriched among upregulated genes by siFOSL2 (Fig 6E). This indicates that FOSL2 may function primarily as a repressive factor in LSCs. Relevant upregulated genes included *TGFBI* and *JAG1* which have known roles in angiogenesis, and *CLDN1* which is involved in tight junctions (Fig 6F). Furthermore, FOSL2 binding signals were identified near these genes (Fig 6G), indicating that these genes are direct targets of FOSL2.

Interestingly, 103 downregulated genes upon si*FOSL2* did not give any enrichment in GO analysis, suggesting a less prominent role of FOSL2 in gene activation. Nevertheless, *CLDN4* and *CLDN7*, other tight junction genes, are significantly downregulated (Fig 6C, F and supplementary Fig 11G), suggesting an altered barrier function in FOSL2-deficient LSCs. Other downregulated genes include *TGM1* and *ABCA12*, linked to epidermal permeability barrier disorders (supplementary Fig 11G).

To summarize, our results demonstrated that, among TFs that define the LSC fate, FOSL2 is a novel LSC TF associated with cornea opacity.

## Discussion

The corneal epithelium and the epidermis are both stratified epithelia, serving as barriers and the first-line defense against external insults. Nevertheless, they have distinct tissue-specific functions that are tightly controlled by the proliferation and differentiation program of their corresponding stem cells, LSCs for the cornea and basal KCs for the epidermis. In this study, we characterized molecular signatures defining the cell fates of these two cell types by integrating in-house multi-omics data as well as publicly available datasets. Using motif and gene regulatory network analyses, we identified a collection of shared and cell type specific epithelial TFs defining KCs and LSCs. Furthermore, we showed a proof-of-principle that this resource of LSC TFs and their regulatory mechanisms can provide novel tools for dissecting pathomechanisms of corneal diseases.

In contrast to the well-studied TFs and their associated gene regulatory networks in epidermal KCs (11–14,57), TFs regulating cornea LSCs have only recently started to emerge. Except for PAX6, identified TFs including p63, SMAD3, RUNX1, and FOXC1 (30,32,39) which are all also expressed in KCs (11–14,57). This raises interesting questions about whether these TFs are sufficient to determine cell fate differences between LSCs and KCs and how they control cell fate determination mechanisms. By specifically comparing LSCs to KCs, we identified PAX6, SMAD3, OTX1, ELF3, and FOSL1 as the LSC specific TFs that determine the LSC fate. To note, these LSC specific TFs may be good candidates for developing (trans)differentiation strategies to generate LSCs from other cell types for corneal regenerations (58,59). The identification of PAX6 as an LSC specific TF was expected. It is an eye development master regulator (30) and is associated with the disease aniridia where corneal opacity is one of the main manifestations (33). Furthermore, PAX6 has previously been shown to co-regulate target genes with RUNX1 and SMAD3 (32). Although SMAD3 is also expressed in KCs, it has higher expression in LSCs, and therefore it is identified to have a higher influence score LSCs than in KCs in our study. OTX1 is an important TF for regulating the neural lineage (60,61). In mice, both Otx1 and its ortholog Otx2 are vital for tissue specification during eye development, particularly of the retinal pigmented epithelium (62,63). ELF3 has previously been linked to *KRT12* and *KRT3* regulation (64), and is one of the TFs identified to play a major role in LSC stratification (39).

FOSL1, along with FOSL2, a novel candidate for corneal disease in our study, and other TFs from the FOS and JUN families, are known to be important in epidermal cells. These TFs can form the AP1 complex together with JUN factors (65), which regulates various biological processes, including epidermal stratification (66). In our differential motif analysis, the FOS motif that can be bound by FOS, FOSL1 and FOSL2, is the most abundant motif enriched in LSCs, as compared to KCs, highlighting the potential role of FOS,FOSL1 and FOSL2 in LSCs.

We found that only a small number of TFs have been identified as KC specific, including GATA3, CEBPA, and HOX genes. Among these TFs, GATA3 and CEBPA have been reported to have a role in the epidermis. CEBPA has been shown to regulate p63 expression (67), while GATA3 is regulated by p63 in the epidermis (68–70). Interestingly, GATA3 was upregulated in aniridia patient LSCs, which is in line with the concept that loss of PAX6 in aniridia LSCs could lead to a KC-like signature. However, our GSEA analyses did not show significant gene expression similarity between upregulated genes in aniridia LSCs and KCs, indicating that aniridia patient LSCs do not acquire a complete KC fate. Surprisingly, we detected HOXA9 as one of the KC specific TFs. HOX genes are well known in antero-posterior body patterning and segmentation where mesodermal genes and cells are mainly involved (71,72), but little is known about their function in the epidermis. One plausible interpretation is that, as LSCs are from the eye and KCs are from the trunk of the body, the detection of HOX9 for the KC fate simply marks the positional information along antero-posterior axis. Nevertheless, the lower number of KC specific TFs, as compared to LSC specific TFs, indicates that repression of LSC specific TF expression is critical for the KC fate. This is in line with our observation that LSC specific TFs such as the *PAX6* locus is completely covered by H3K27me3, probably via polycomb repression. It is also worth noting that the ANANSE prediction tool used in this study is unable to reliably predict TFs with transcriptional repression functions (52), which limits the identification of TFs to repress LSC genes, if there are any in KCs. Consistent with this, ANANSE did not detect repressive TFs such as OVOL2 (73) and SOX9 (74) that have been shown to play roles in LSCs.

We anticipated that TFs that are shared but important to both LSCs and KCs could be not identified through pairwise gene regulatory network comparison between LSCs and KCs. For detecting these shared TFs, we compared both cell types to pluripotent stem cells in the gene regulatory network analysis. This approach indeed resulted in a significant number of shared epithelial TFs including FOSL2, p63, EHF, TFAP2A, KLF4/5, FOS, JUN, RUNX1 and FOXC1. Many of these have previously been linked to important functions in both epidermis and cornea (12–14,16,32,75–79).

The key role of TFs in cell fate control is often demonstrated by their association with developmental diseases. PAX6, FOXC1 and p63 are known to be associated with corneal opacity (20,33,37), and FOSL2 is a novel corneal opacity gene identified in this study. Except for PAX6, both FOXC1 and FOSL2 are shared between LSCs and KCs. FOSL2, PAX6 and FOXC1 seem to regulate most identified disease genes. For PAX6 and FOXC1, this is consistent with their broad expression patterns in the eye and the phenotypic heterogeneity and overlap linked to *FOXC1* and *PAX6* mutations, e.g., iris and corneal defects and higher prevalence of glaucoma (33,37), reinforcing a common regulatory network shared by the two TFs (80). As for PAX6 and p63 associated disorders, although *PAX6* and *TP63* mutations are known to cause corneal opacity, other phenotypes are quite distinct, fully in line with their gene expression in different tissues, e.g., *PAX6* in the cornea epithelium, iris, retina, pancreas and parts of the central nervous system, and *TP63* in the cornea epithelium, skin epidermis and other stratified epithelia (20,33). Although FOXC1 was annotated as a shared TF of LSCs and KCs in our study, its relevance for the skin and cornea might still be different. In our analysis, FOXC1 had a higher influence score in LSCs, probably due to its higher expression in LSCs. This is in line with skin phenotypes not being reported in anterior segment dysgenesis associated with *FOXC1* mutations.

In our analysis, we identified two TFs that are part of the AP1 complex. FOSL1, which is mainly an activating factor (81), was identified as an LSC specific TF. In line with this, decreased expression of FOSL1 in the cornea has been linked to keratoconus patients (82). Next to FOSL1, we also identified FOSL2 as a shared epithelial TF and a novel TF associated with corneal opacity, which was corroborated by *in vitro* and *in vivo* protein expression. Furthermore, we identified FOSL2 as a direct target gene of PAX6, and it was downregulated in PAX6-deficient cells, enforcing the role of FOSL2 in corneal pathology. In mice, Fosl2 has been linked to extensive dermatosis (81). and abnormalities in the cornea and anterior segment, but these were mainly overexpression studies (83), whilst in humans *FOSL2* has been linked to skin, with a recent GWAS study linking SNP-associated with increased FOSL2 expression to a higher likelihood of eczema (84). More importantly, *FOSL2* truncating variants have recently been linked to a neurodevelopmental syndrome with multiple ectodermal symptoms including scalp aplasia cutis (absence of scalp skin) and tooth and nail abnormalities (85). Patients also presented with congenital cataracts, but no corneal phenotype was discovered; however, these often manifest at an older age than the patients’ age reported in the study.

In this study, we have found a predicted-pathogenic heterozygous missense variant in *FOSL2* in a proband with corneal opacity. Upon siRNA-mediated knockdown in LSCs, we observed that reduced FOSL2 led to an upregulation of pro-angiogenesis genes like *JAG1* and *TGFBI.* Furthermore FOSL2 binding sites were also found in their associated loci, indicating a role of FOSL2 in repressing angiogenesis. Corneal neovascularization is one of the hallmarks of LSCD and corneal opacities. FOSL2 has been identified to promote VEGF independent angiogenesis in fibroblasts (86), and *FOSL2* knock down in glioma endothelial cells increased blood tumor barrier permeability (87). The latter study also showed that FOSL2 binds to the promoters and downregulates the expression of several tight junction genes *TJP1*, *OCLN* and *CLDN5*, increasing barrier permeability. In accordance, we also detected altered expression of many tight junction and barrier genes in our siFOSL2 LSCs, such as *CLDN1*, *CLDN4* or *TGM1*. Tight junctions are essential for the cell-cell contact and permeability barrier function in epithelial cells, and defects in tight junctions are linked to several epithelial diseases (88,89). Therefore, deregulation of tight junction genes and upregulation of pro-angiogenic genes upon *FOSL2* knockdown strongly support FOSL2 relevance in LSCs identity and the corneal opacity phenotype.

In this study, we identified shared and cell type specific epithelial TFs that are important in determining cell fate of LSCs and KCs. Furthermore, we provided evidence for potential TFs and regulatory mechanisms in corneal diseases, by uncovering a novel role for FOSL2 in corneal opacity. We therefore show proof of principle that these key LSC TFs and their gene regulatory networks can be leveraged as a resource to unveil pathomechanisms behind corneal opacity diseases.

## Data availability

All non-patient cell raw sequencing files generated in this study have been deposited in the GEO database with the accession number GSE206924. All aniridia-patient sequencing files have been deposited in the EGA database with controlled access with the study number EGAS00001007397. Publicly accessible data was downloaded from GEO using the accession codes provided in supplementary tables 1, 2, and 3. There are no restrictions on data availability. Source data are provided with this paper All code used in this study is available at https://github.com/JGASmits/regulatory-networks-in-epidermal-and-corneal-epithelia All epigenetic data is available in a UCSC genomebrowser trackhub via: https://mbdata.science.ru.nl/jsmits/epi_fate/epi_fate_trackhub.hub.txt

## Abbreviations

CRE: Cis-regulatory element
ESCs: Embryonic stem cells
FDR: False Discovery Rate
FECD: Fuchs Endothelial Corneal Dystrophy
GO: Gene Onthology
GSEA: Gene set enrichment analysis
KCs: Keratinocytes
LSCD: Limbal stem cell deficiency
LSCs: Limbal stem cells
scRNA-seq: single-cell RNA-seq
TF: Transcription factor

## Author contribution

JGAS, SJH, and HZ designed the research; JGAS, DLC, MA, MB, CL, JQ, TS, LL and LNR performed experiments; JGAS, SJH, and HZ analyzed multi-omics data; MA, TS, MB, LL, LNR, NS, BS and DAcontributed and performed limbal stem cell culture experiments, NO and MM analyzed whole genome sequence data; MA, TS performed siRNAseq experiments, MB performed immunostaining experiments, JGAS, SJH and HZ wrote the original draft; all authors edited and contributed to the manuscript.

## Acknowledgement

We thank M.P.A. Baltissen, L.A. Lamers and S. Rinzema for operating the Illumina analyzer and initial data demultiplexing, L. Wingens for support with the celseq library preparations, J. Arts, and W.N. Twilhaar for their pre-processing of the single cell RNAseq objects of the in vivo datasets, J. Niehaus for support in generating the FOSL2 CUT&RUN tracks. This work was supported by the Aard-en Levenswetenschappen, Nederlandse Organisatie voor Wetenschappelijk Onderzoek NWO-ALW (ALWOP 376 to JGAS, HZ), the European Joint Programme Rare diseases EJP-RD/ZonMw (JPRD20-135/463003005 to DLC, HZ), and by the European Cooperation in Science and Technology COST (CA18116 to DLC, LL, SF, NS, DA, HZ). Among these fundings, JGAS received the salary from NWO-ALW (ALWOP 376) and DLC from EJP-RD/ZonMw (JPRD20-135/463003005). The funders had no role in study design, data collection and analysis, decision to publish, or preparation of the manuscript.

This research was made possible through access to the data and findings generated by the 100,000 Genomes Project. The 100,000 Genomes Project is managed by Genomics England Limited (a wholly owned company of the Department of Health and Social Care). The 100,000 Genomes Project is funded by the National Institute for Health Research and NHS England. The Wellcome Trust, Cancer Research UK and the Medical Research Council have also funded research infrastructure. The 100,000 Genomes Project uses data provided by patients and collected by the National Health Service as part of their care and support.

## Materials and Methods

### Ethical statement

All procedures for establishing and maintaining human primary keratinocytes were approved by the ethical committee of the Radboud university medical center (“Commissie Mensgebonden Onderzoek Arnhem-Nijmegen”) (CMO-nr:2004/132). All donors have given informed consent in written forms.

All procedures for establishing and maintaining limbal stem cells were conducted according to the principles expressed in the Declaration of Helsinki. The use of limbal stem cells extracted from corneal scleral donor rims was approved by the Ethics Committee of the Saarland (Number 226/15 and 110/17). Aniridia patients consented in written form to research limbal biopsy during ocular surgery.

Permission to use the donor corneas for research purposes was given by the donor’s next of kin by signing a written consent form released by the Regional Transplant Service and the National Transplant Service, without ethics committee approvals. According to Article 4 of the Italian Law 91 of 1st April 1999, donor corneas that cannot be used for transplantation (i.e., unsuitable for biological reasons or anamnestic reasons) can be used for research purposes if the aim is to ameliorate corneal transplantation or progress towards a cure of corneal diseases. The human corneas used in this study were unsuitable for transplantation and obtained by Fondazione Banca degli Occhi del Veneto (www.fbov.org, Venice, Italy) for research purposes.

### KC and LSC cell culture in vitro

KCs were isolated and cultured as previously described(16). Briefly, after isolation primary KCs were cultured in Keratinocyte Basal Medium supplemented with 100 U/mL Penicillin/Streptomycin, 0.1 mM ethanolamine, 0.1 mM O-phosphoethanolamine, 0.4% (vol/vol) bovine pituitary extract, 0.5 μg/mL hydrocortisone, 5 μg/mL insulin and 10 ng/mL epidermal growth factor. Medium was refreshed every other day until the cells were 90% confluent.

Limbal tissues were acquired as previously described (90). Two aniridia limbal tissue single biopsies were obtained from the superior limbus during penetrating keratoplasty from 2 patients with congenital aniridia as previously described (91). Genetics of the aniridia patients were identified to be c.33delC p.Gly12Valfs*19 (NM_000280.2) for AN55 and c.990_993dup p.Met332Alafs*10 for AN40 (supplementary table 1). Both patients their corneas were in Lagali Stage 4 during the Keratoplasty(31).

Cell isolation was performed as previously described(90). Briefly, limbal tissue was digested in collagenase A solution (4 mg/ml) in keratinocyte serum-free medium (KSFM) (Thermo Fisher Scientific; Waltham, MA) for 20 h at 37 °C. Cell suspensions were filtered through a use of Flowmi® micro strainer (SP Bel-Art; Wayne, NJ). LSC clusters were dissociated with trypsin-EDTA (0.05%) solution and cultivated in KSFM. Medium was refreshed every other day. Subconfluent (80–90%) limbal epithelial cells were harvested at passage 3. Aniridia LSCs were harvested at passage 3.

Next to this approach, other LSC samples (LSC-Aberdam, supplementary table 1) were isolated from postmortem donated peripheral corneal epithelium and cultured as previously described(50). Briefly after isolation, they were expanded and cultured in KSFM (Gibco, Life Technologies) supplemented with 25 μg/ml Bovine Pituitary Extract (BPE; Gibco, Life Technologies), 0.2 ng/ml Epidermal Growth Factor (EGF, Peprotech, Neuilly-sur-Seine, France), 0.4 mM CaCl_2_, 2 mM Glutamine (Gibco, Life Technologies) and 100 U/ml Penicillin/Streptomicin (Gibco, Life Technologies). Medium was refreshed every other day until the cells were 90% confluent.

Finally for CUT&RUN of LSCs, LSCs were isolated and cultured as described previously (see paper by Fasolo et al and by Luznik et al (92,93). Briefly, corneal limbal tissue was dissected from human donor corneas, cut into small pieces, and incubated with 10ml trypsin (0.05% trypsin-0.01% EDTA solution, Life Technologies, 25300-062) for 30 minutes at 37°C for 4 consecutive times. Every time, the supernatants containing the cells were collected, neutralized, centrifuged at 1000rpm for 5 minutes and cells co-cultured with a feeder layer (consisting of lethally irradiated 3T3-J2 cells) in LSC growth medium. The LSC growth medium contains 2:1 Dulbecco’s Modified Eagle Medium (Life Technologies, 21969035) and F12 (Life Technologies, 21765029), 10% Fetal Bovine Serum (Life Technologies, 10099-141), 50 mg/mL penicillin-streptomycin (Life Technologies, 15140122), 4 mM glutamine (Life Technologies, 25030081), 0.18 mM adenine grade I (Pharma Waldhof GMBH, 4010-21-2), 0.4 mg/mL hydrocortisone (Flebocortid Richter, Sanofi, AIC013986029), insulin (Humulin R, Lilly, Canada, HI0210), 2 nM triiodothyronine (Liotir, IBSA, AIC036906016) and 8.1 mg/mL cholera toxin QD (List Biological Laboratories, 9100B). Cultures were incubated at 37°C with 5% CO_2_. Medium was changed every 2 days with further addition of 10 ng/mL EGF (Cell Genix GmbH, Germany 1416-050). Cultures were switched to KSFM and used after three passages

### siRNAseq treatment

Primary human limbal epithelial stem cells were isolated from healthy donors and cultivated in 6 well plates in KSFM medium (Cat. Nr. 17005042, Thermo Fisher, Gibco, Life Technologies, Paisley, UK) supplemented with 500 ng Epidermal growth factor (EGF), 12.5 mg Bovin pituitary extract (BPE) (Cat. Nr. 37000015, Thermo Fisher, Gibco, Life Technologies, Paisley, UK) and 0.1% P/S. After reaching 70-80 % confluence, cells were transfected using Lipofectamin 2000 transfection reagent. For this purpose, 5 l Lipofectamin was added to 250 l Optimem and incubated for 5 minutes, at room temperature. In a separate tube, siRNA of interest (information and concentrations used are summarized in Table 1) was diluted in 250 l Optimem. The Lipofectamin was added to the siRNA mixture and incubated for 20 minutes, RT. After the incubation time, transfection mixture was added dropwise to the cells. The cells were incubated at 37°C and 5% CO2 and medium were changed after 24h. Cells were collected 48 hours after transfection for further analysis.

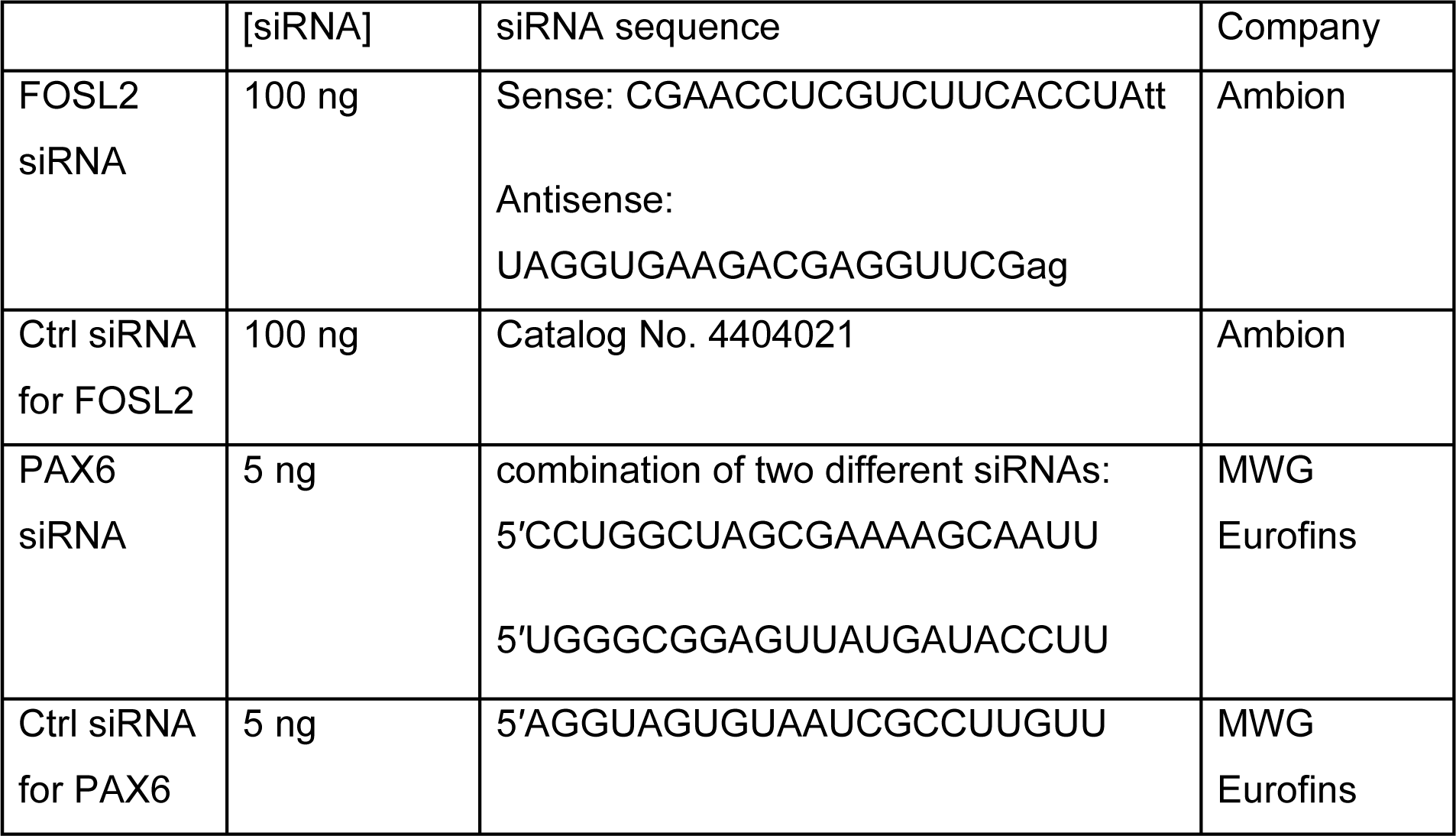

### Bulk RNA-seq

Total RNA was isolated using the Quick–RNA MicroPrep kit (Zymo Research), according to the manufacturer’s protocol. RNA concentrations were measured using the the DeNovix DS-11FX spectrometer. 500 ng of RNA was prepared for sequencing using the KAPA RNA HyperPrep Kit with RiboErase (Kapa Biosystems). Libraries were sequenced on the NextSeq 500 (Illumina), generating an average of 15–20 million reads per sample.

### Single-cell RNA-seq library preparation and sequencing

A single-cell suspension was made using trypsin. After which cells were filtered using a 40uM filter to remove cell clumps. Cells were stained with 7-AAD. The live cells were selected for and FACS-sorted onto 384-well plates containing primers with unique molecular identifiers, according to the SORT-Seq protocol(94). Plates were spun down (1200 × *g*, 1 min, 4 °C) and ERCC spike-in mix (1:50,000) was dispensed by a Nanodrop (BioNex Inc) into each well. 150 nl of the Reverse Transcription (RT) mix was dispensed into each well. Thermal cycling conditions were set at 4 °C 5 min; 25 °C 10 min; 42 °C 1 h; 70 °C 10 min. The library of each plate was pooled together and the cDNA was purified using AmpureXP (New England BioLabs) beads. Overnight in vitro transcription (Ambion MEGA-Script) was carried out at 16 °C, with the lid set at 70 °C. An exonuclease digestion step was performed thereafter for 20 min at 37 °C, followed by fragmentation of the RNA samples. After a beads cleanup, the samples were subjected to library RT and amplification to tag the RNA molecules with specific and unique sample indexes (Illumina), followed by a final beads cleanup (1:0.8, reaction mix: beads) and the sample cDNA libraries were eluted with DNAse free water. Libraries were quantified using the KAPPA quantification kit following manufacturers protocol after which the plates were sequenced on the NextSeq 500 (Illumina) for 25 million reads per plate.

### ChIP-seq

Chromatin for ChIP was prepared as previously described(95) with minor modifications. On average, 0.5M cells were used in each ChIP. Antibodies against H3K27ac (Diagenode #C15410174, 1.2 μg), H3K4me3 (Diagenode #C15410003, 1 μg), H3K27me3 (Diagenode #C15410069, 1.5 μg), p63 (Santa Cruz #H129, 1 μg, recognizing the C-terminal α tail of p63) were used in ChIP assay. Afterwards 5ng DNA fragments were pooled and proceeded on with library construction using KAPA Hyper Prep Kit (Kapa Biosystems #KK8504) according to the standard protocol. The prepared libraries were then sequenced using the NextSeq 500 (Illumina) according to standard Illumina protocols.

### Immunofluorescence stainings

Human donor corneas were fixed with 4% paraformaldehyde overnight at 4°C, soaked in increasing gradients of sucrose solutions (7.5%, 15% and 30% in PBS, at least 30 min each at 4°C, the last one overnight) and then embedded in OCT compound, frozen, and cut into 5- to 7-mm sections. Sections were permeabilized in a solution containing 0.5% Triton X-100 for 10 minutes, incubated in 5% BSA (Sigma-Aldrich A7906) for 1 hour at room temperature and then incubated overnight at 4°C with primary antibodies. The following primary antibodies (all diluted at 1:100) were tested: PAX6 (rabbit, Biolegend Poly19013); FOSL1 aka Fra-1 (C-12) (mouse, Santa Cruz Biotechnology, sc-28310); Anti-FOSL2 (rabbit, Sigma, HPA004817); p63 (mouse, DAKO, M7317). Sections were extensively washed in 1X PBS and incubated with secondary antibodies (Alexa Fluor 488 anti-rabbit A11008, Rhodamine Red-X goat anti-mouse IgG R6393, Rhodamine Red-X goat anti-rabbit IgG R6394, Alexa Fluor 488 goat anti-mouse A11001, all from Invitrogen, all diluted at 1:500) for 45 minutes at room temperature. Sections were eventually mounted with medium containing DAPI Fluoromont-G (EMS, Società Italiana Chimici, Rome, Italy, 17984-24) and pictures taken and evaluated through an Eclipse Ti Nikon microscope (Nikon, Amstelveen, The Netherlands).

LSCs and primary keratinocytes were grown in cell culture chamber slides (Corning, NY USA), and, when confluent, fixed using 4% PFA solution. Immunostainings were performed as previously described(96). Briefly, cells were washed, permeabilized for 15min with 0.1% Triton X-100, washed again and blocked for 2h in 1% FBS. Primary antibody incubation was done overnight at 4οC with the following antibodies diluted in blocking buffer: PAX6 (1:300, 901301 Biolegend), P63 (1:100, sc8344 SCBT), FOSL1 (1:100, sc20310, SCBT), FOSL2 (1:200, HPA004817 Sigma Aldrich). Cells were washed thoroughly and incubated with secondary antibodies AlexaFlour 488 (1:400, A11008 Invitrogen) and AlexaFluor 647 (1:400, A31571 Invitrogen, CA, USA). Slides were mounted with Vectashield (VectorLabs, CA, USA) and analyzed with Zeiss AI Sample Finder (Zeiss, Jena, Germany). Fiji/ImageJ software (National Institutes of Health, MD, USA) was used for overlay images preparation.

### CUT&RUN

CUT&RUN was performed using the CUT&RUN Assay Kit (Cell signaling Technology #86652) and according to the manufacturer instructions. Briefly, 200 000 live primary LSCs were used per immunoprecipitation (IP). For FOSL2 IPs, 10mL of antibody (D2F1E, Rabbit mAb #19967) was used per IPs, two IPs were performed per replica to ensure a sufficient amount of DNA. The negative control IgG included in the kit was used to check for enrichment by quantitative PCR.

For high-throughput sequencing, libraries were prepared using KAPA HyperPrep Kit (Kapa Biosystems #KK8504) using 5ng of DNA. Libraries concentration and fragment repartition were checked by automated electrophoresis (Bioanalyzer Instrument, Agilent). Libraries were sequenced using the NextSeq 500 (Illumina) and according to their protocol.

### Reverse transcription quantitative PCR (RT-qPCR) validation

### Disease target genes

250ng RNA were used for cDNA synthesis using the iScript cDNA synthesis kit (BioRad, CA, USA). Quantitative PCR (qPCR) was carried out using 4 μl cDNA (diluted 1:20) with iQ SYBR Green Supermix (BioRad, CA, USA) and specific primers (Biolegio, Nijmegen, NL):

*GAPDH* Forward (Fw): aag gag taa gac ccc tgg acc a,

*GAPDH* Reverse (Rv): gca act gtg agg agg gga gatt

*ACTB* Fw: TTC TAC AAT GAG CTG CGT G

*ACTB* Rv: GGG GTG TTG AAG GTC TCA AA

*JAG1* Fw: GAT CGC CTG CTC AAA GGT CT

*JAG1* Rv: GAC TGG AAG ACC GAC ACT CG

*TGFBI* Fw: CCT TTG AGA CCC TTC GGG CTG

*TGFBI* Rv: TCG AAG GCC TCA TTG GTC GG

*CLDN1* Fw: CCC CAG TCA ATG CCA GGT ACG

*CLDN1* Rv: TCG GGG ACA GGA ACA GCA AA

*OCLN* Fw: TCT AGG ACG CAG CAG ATT GGT

*OCLN* Rv: TCA GGC CTG TAA GGA GGT GG

All disease gene qPCRs were performed using a BioRad CFX96^TM^ system following standard cycling conditions. Data were analyzed by the ΔΔCT method, normalized to housekeepers *GAPDH* and *ACTB*.

### FOSL2 and PAX6 siRNA

RNA extraction were performed using RNA isolation kit (RNA Purification Plus Micro Kit, Cat. Nr. 47700, (Norgen Biotek CORP. Canada)), according to the manufacturer’s instructions. Optional DNA column digestion twas performed. The eluted RNA and protein samples were stored at −80°C until further use. NEB One Taq RT-PCR kit (One Taq® RT-PCR Kit, New England Biolabs INC, Frankfurt, Germany) was used for cDNA synthesis, following the manufacturer’s instructions. Samples were stored at -20 °C. The quantitative Polymerase Chain Reaction (qPCR) mix contained 1 µl of the specific primer solution, 5 µl SYBR Green Mix (Qiagen N.V., Venlo, Netherlands), and 3 µl nuclease-free water (total volume: 9 µl). To investigate the mRNA expression levels, 1 µl cDNA of interest was added to the qPCR mix. The “QuantStudio 5 real-time PCR system” (Thermo Fisher Scientific, Waltham, MA, USA) was used for the qPCR reactions. The amplification conditions were 95°C for 10s, 60°C for 30s, and 95°C for 15s (40 cycles). Samples were measured in duplicates and experiments were repeated for at least 5 donors. Values were normalized to the expression value of TATA-Box binding protein as an endogenous control gene using the ΔΔCT method and the fold change (2ΔΔCT-value) was used for statistical analysis. All primers were purchased from Quantitect Primer assay ((Qiagen N.V., Venlo, Netherlands; PAX6: QT00071169, FOSL2: QT01000881 and TBP: QT00000721).

### RNA-seq, ATAC-seq ChIP-seq and CUT&RUN data preprocessing

Preprocessing of reads was done automatically with workflow tool seq2science v0.7.1(97). Paired-end reads were trimmed with fastp v0.20.1(98) with default options. Genome assembly GRCh38.p13 was downloaded with genomepy 0.11.1(99). Public samples were downloaded from the Sequence Read Archive(100) with help of the NCBI e-utilities and pysradb(101). The effective genome size was estimated per sample by khmer v2.0(102) by calculating the number of unique kmers with k being the average read length per sample. scATAC fastq files were merged to pseudobulk by combining all fastq files from each plate using the bash command cat.

Reads of ChIP-seq, CUT&RUN and ATACseq were aligned with bwa-mem v0.7.17(103) with options ‘-M’.

Reads of RNAseq samples were aligned with(104) with default options. Afterwards, duplicate reads were marked with Picard MarkDuplicates v2.23.8(105). General alignment statistics were collected by samtools stats v1.14(106). Mapped reads were removed if they did not have a minimum mapping quality of 30, were a (secondary) multimapper or aligned inside the ENCODE blacklist(107). RNAseq sample counting and summarizing to gene-level was performed on filtered bam using(108). Sample sequencing strandness was inferred using(109) in order to improve quantification accuracy.

ATAC samples were tn5 bias shifted by seq2science. ChiP, CUT&RUN and ATAC sample peaks were called with macs2 v2.2.7(110) with options ‘--shift -100 --extsize 200 -- nomodel --keep-dup 1 --buffer-size 10000’ in BAM mode. The effective genome size was estimated by taking the number of unique kmers in the assembly of the same length as the average read length for each sample. Narrowpeak files of ChiP-seq and CUT&RUN biological replicates belonging to the same condition were merged with the irreproducible discovery rate v2.0.4.2(111). ATAC-seq samples were correlated seq2science its DESeq2 reads per peak spearman correlation clustering (supplementary Fig 5F).

### Single-cell RNA-seq data preprocessing

Single-cell libraries were pre-processed using the cellseq2 pipeline. Briefly, reads were aligned using star to the GRCh38.p13 genome. After which cells were quality controlled using Seurat, filtering cells on ERCC reads, genes measured and transcripts per cell. After visualization of the lack of heterogeneity by Umap, pseudobulk count data was generated by summing all the cells their UMI counts. Cellular heterogeneity was assessed using the analysis file Generate_scRNAseq_pseudobulk.Rmd. Finally single-cell and bulk gene count tables were merged for a combined bulk and pseudobulk analysis.

### RNA-seq data analysis and normalization

The bulk and pseudobulk count tables were merged on gene names, keeping all gene names detected. Due to potential sex differences between donors, genes located on chromosome X and Y were removed. Finally, genes with less than 10 counts per row were removed. Variance visualization was performed using sample distance and PCA. For quality control, sample variance and distance were visualized before and after removing technical variance due to different sequencing methods. Limma (112) was used to remove these batch effects.

Rld normalization was used for normalizing gene intensities. Between all conditions differentially expressed genes were detected using Deseq2 (113). Non-batch corrected count tables were used for identifying the differentially regulated genes. Ashr log2 fold change shrinkage (114) was used to shrink the Log2 fold change values. Differentially regulated gene detection cutoffs were set as an adjusted *p* value of 0.01 or lower, and an absolute log2 FC of 0.58 and larger.

The packages complex Heatmaps (115) and circlize (116), were used to visualize the differentially expressed genes. Subsequently Progeny enrichment (47) was performed to quantify signaling pathway target gene enrichment. Clusterprofiler (44) was run for GO-term enrichment on differentially expressed genes of each comparison. Finally, foldchange of all genes were used to generate a gene list for GSEA enrichment of the MSigDB collections (46). Gene names were mapped to ENTREZID using AnnotationDbi (117) and these were used to run KEGG pathway enrichment. The enriched pathways were visualized using pathview (118).

### Identification of CREs

In order to identify CREs, ATAC-seq was used. Bulk and scATAC data were merged from *in vitro* expanded KCs and LSCs. Next to the generated datasets, publicly available data were incorporated. To prevent a sequencing depth bias, the top 100.000 ATAC peaks from each cell type were combined, the overlapping peak summits were merged, and histone modifications in varying window sizes around these ATAC peaks were quantified using histone ChIP-seq datasets (Fig 2A). For ATAC signal quantification the ATAC intensity was quantified in 200bp around the peak summits, and for the promoter mark H3K4me3 and the enhancer mark H3K27ac a 2kb window was used and finally for the repressive H3K27me3 mark a 5kb window was used for quantification (Fig 2A). This resulted in an extensive dataset containing cis regulatory elements and their respective histone modification signal intensity.

Differential CREs were identified for the ATAC-seq and H3K27ac reads by running DESEQ2 on the read counts within the defined windows and identified regions (adjusted p-value < 0.05). For H3K4me3 and H3K27me3 signals, Differential CREs were identified with two steps. First the histone mark distribution was plotted, and CREs with a low to no histone signal were disregarded. Next, high activity regions with variable signal were selected. (supplementary Fig 5).

Variable cis-regulatory elements were linked to genes with two approaches

1. CREs were linked to all the TSS regions within a 100kb window using bedtool window(119) after which the CREs per TF were distance weighted and summed based on the ANANSE distance weighing approach including promoter peaks(52).
2. CREs were linked to the closest TSS region within 20kb using bedtool closest(119).

After linking the regions to genes in both approaches, intensity scores were printed to a CSV file. Heatmaps and go term enrichments were generated in R using clusterprofiler and complex heatmap.

### Single-cell ATAC-seq

A single-cell suspension was made using trypsin. After which cells were filtered using a 40uM filter to remove cell clumps. The protocol on from Chen et all(120) was used to sequence single-cell ATAC. Briefly 50.000 cells were tagmented in bulk in 45 μl of tagmentation mix (20mM Tris pH 7.6, 10mM magnesium Chloride 20% Dimethylformamide), 5μl of tagmentation protein and 0.25 μl of Digitonin. Cells were tagmented for 30min at 37 °C and 800 rpm.

Tagmentation was stopped by adding 50 μl of tagmentation stop buffer (10mM Tris-HCL PH 7.8 and 20 mM EDTA). Cells were stained with DAPI and DAPI positive cells were FACs sorted in 384-well plates containing Nextera primers with unique molecular identifiers, NACL ProteinaseK and SDS page. Plates were spun down (1200 × *g*, 1 min, 4 °C) and were incubated for 15min at 65 °C.

4μl of Tween20 was added. 2μl of H20 was added and finally 10 μl of NEBNext High-Fidelity 2X PCR Master Mix was added to each well. Thermal cycling conditions were set at 72 °C 5 min; 98 °C 5 min; and then 20 repeats of 98°C for 10s, 63°C for 30s, 72°C for 20s.

Plate libraries were pooled and purified using a Qiagen PCR purification kit with adjusted buffer volumes according to Chen et all(120). After the column cleanup a final beads cleanup was performed using AmpureXP (New England BioLabs) beads and the sample cDNA libraries were eluted with DNAse free water. Libraries were quantified using the KAPPA quantification kit following manufacturers protocol after which the plates were sequenced on the NextSeq 500 (Illumina) for 30 million reads per plate.

### Motif analysis

The Gimme motifs database was pre-filtered(51) to include only motifs linked to TFs which were expressed in either KC and/or LSCs (using a cutoff of at least 10 counts in total). When multiple motifs mapped to a TF, the most variable motif was used. In case multiple TFs mapped to a motif, the most differential TF on the transcriptome was annotated to the motif. All highly variable CREs their log10 quantile normalized values were used as an input for Gimme maelstrom motif enrichment analysis.

### ANANSE analysis

For the gene regulatory network analysis, all the called ATAC peaks were used, merging summits and excluding peaks on the chromosomes GL, Un, KI, MT, X, and Y due to potential donor sex differences. Next, ANANSE binding was ran using all the peaks as potential enhancer regions and using both ATAC and H3K27ac signals to predict potential TF binding. To select the TF binding model a Jaccard similarity score of min 0.2 was used, to minimalize the false-positive models used to predict TF binding. For ANANSE network, the ANANSE binding files were combined with the RNAseq TPM files. This included all bulk RNAseq samples of KCs, LSCs and ESCs (supplementary Table 1). In the case of the *in vivo* pseudobulk data, FPKM values were used based on the UMI tables.

Finally, ANANSE influence was ran using the top 500.000 differential edges between networks. Deseq2 was ran on the countfiles of each comparison to identify differential genes needed for ANANSE influence. To prevent missing values, for the final ESC-KC, ESC-LSC, KC-LSC and LSC-KC comparison all differential edges were taken from each comparison and used to reran each comparison with all these edges included. This prevented missing values in the differential networks while comparing different differential networks.

TF hierarchy was estimated using the TF-target TF binding score generated by the influence command running the –full-output flag. This represents the motif, ATAC&H3K27ac signal intensity in the target TF locus and is excluding the difference in expression. The Delta binding score was calculated by subtracting the score of a TF-gene interaction within one GRN with the score of the interaction within the other GRN.

The delta binding score of the ESC-KC and ESC-LSC comparisons were averaged. If this average was higher than the delta binding score of KC-LSC and LSC-KC the interaction was classified as ‘shared epithelial’, if the delta binding score was highest in LSC-KC it was classified as ‘KC specific’, if the delta binding score was highest in KC-LSC it was classified as ‘LSC specific’.

### Single-cell RNA-seq analysis of the epidermis and the cornea

The raw sequencing data was downloaded from GEO and split it into fastq files using seq2science. Cellranger count was run with Cellranger 6.0.1 to retrieve the matrix, barcodes, and features files necessary for Seurat(121) analysis in R. scRNA-seq cells were selected with a minimum count of 2000, a feature number above 1000, and a mitochondrial percentage below 30 percent. Cell cycle scoring was performed using Seurat CellCycleScoring() feature with the cell cycle genes from Tirosh et al(122). All cells not in the G1 phase were removed. Leiden Clustering was performed and cell clusters were annotated based on described marker genes.

For the data of the epidermis, cell clusters were selected with high KRT14, KRT5, and low KRT1 and KRT10 expression as basal KCs. From the cornea dataset, cell clusters with high S100A2 with PAX6 and TP63 expression and without CPVL expression were selected as LSCs. The *in vivo* vs *in vitro* fold change difference plot was generated by loading deseq2 result tables to identify the TF fold changes.

### ChIP-seq and CUT&RUN data analysis

ChIP-seq and CUT&RUN peaks were called with MACs2 and validated by IDR (preprocessing). Next for each peak summit reads were counted in 200bp windows across each summit. Values were log-transformed and quantile-based normalized. Peaks were linked to TSS regions in 100kb, using bedtools window. Afterwards they were distance weighted using the ANANSE distance weighing approach. When genes did not have ChIP-seq peaks within a 100kb window, they got an intensity score of 0.

Disease gene lists were collected from the EyeDiseases database(56), including all disease gene lists of more than 20 genes. A one-sided Man-Whitney U test was performed to test the hypothesis that the disease genes have more TF binding than the other genes in the genome. Of each significant hit, the top 5 of most bound gene loci were outputted to a list. And the final list was used to generate a dotplot in R (hipseq_intensity_npeak_dotplot.Rmd). Alternatively, ChIP-seq/CUT&RUN peaks were mapped to the gene TSS start site using bedtool closest. Next disease genes mapped vs non-mapped disease genes were compared to all genes mapped vs non-mapped. Using a Fisher exact test.

### Cornea disease gene list

Curated cornea disease list was firstly compiled by retrieving all known genetic disorders affecting the cornea and respective affected genes from “Ophthalmic Genetic Diseases” (123) and then confirmed using available literature in Pubmed (https://pubmed.ncbi.nlm.nih.gov/) and online eye disease (https://gene.vision/) databases. Diseases were grouped as 1) corneal diseases; 2) systemic (or other ocular) disorders with corneal phenotypes, and 3) Diseases with secondary cornea involvement (due to exposure, or unclear involvement). Genes associated with multifactorial disease keratoconus were added based on literature search mainly of published genome wide association (GWAS) and linkage (GWLS) studies(123–125) Curated gene list is available as supplementary Table 7.

### Variant discovery

Participants of the 100,000 Genomes Project were identified for our analyses who had at least one of the following HPO terms or daughter terms present: corneal opacity (HP:0007957), corneal scarring (HP:0000559), Opacification of the corneal stroma (HP:0007759), central opacification of the cornea (HP:0011493), band keratopathy (HP:0000585), central posterior corneal opacity (HP:0008511), corneal crystals (HP:0000531), generalized opacification of the cornea (HP:0011494), peripheral opacification of the cornea (HP:0008011), punctate opacification of the cornea (HP:0007856) and sclerocornea (HP:0000647). A total of 33 probands were identified who remain genetically unsolved. The whole genome sequence data was interrogated for single-nucleotide variants (SNVs) and indels (insertions or deletions), copy number variants (CNVs) and structural variants as previously described (Owen et al 2022). Filtered variants were annotated using Ensembl Variant Effect Predictor (VEP v99) and prioritised identified variants using scores available from CADD, MutationTaster, Provean, Sift, polyphen2, MetaRNN, DANN, fathmm-MKL. Variant nomenclature was assessed using Variant Validator.

---

**Supplementary Fig 1.**
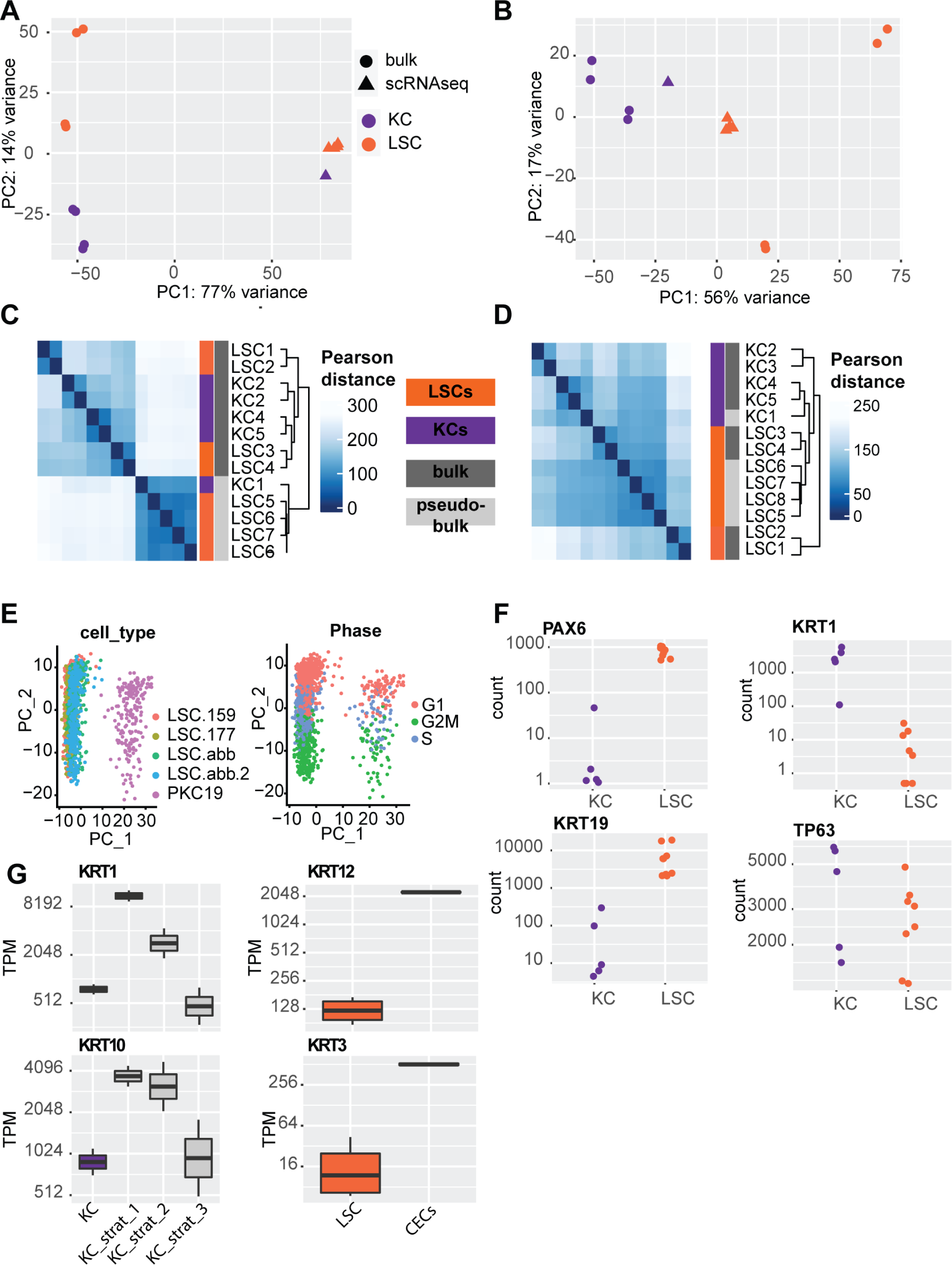
Additional quality control RNA-seq analysis of LSCs and KCs. (A) PCA plot of RNA-seq samples before batch correction. (B) PCA plot after batch correction. (C) Pearson correlation matrix before batch correction. (D) Pearson correlation matrix after batch correction. (E) Umap dimensionality reduction of scRNA-seq data, visualizing the samples each cell is from on the left, and the cell cycle state on the right. (F) Gene count plot for PAX6, KRT1, KRT19, and TP63 in all KC and LSC samples. (G) TPM gene plots for KRT1, KRT10, in KC and various stratified KC samples ranging from day 2 (KC_strat_1), day 4 (KC_strat_2) and day 7 (KC_strat_3) ofratification, and of KRT12 and KRT3 in LSC and air lifted stratified cornea epithelial cells (CECs).

**Supplementary Fig 2.**
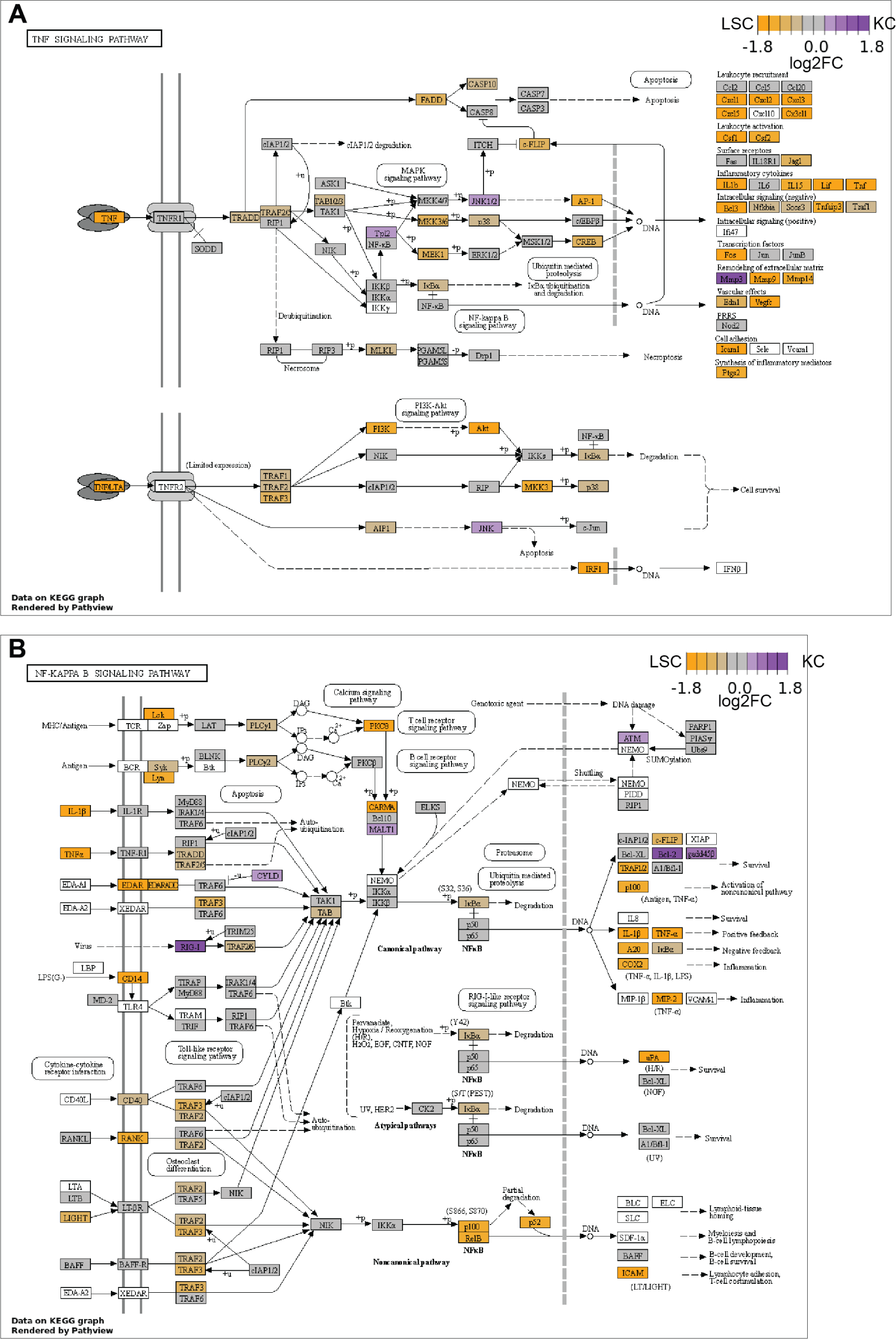
KEGG pathway expression visualizations LSCs and KCs. (A) TNF signaling pathway component expression FC differences between KC and LSCs. (B) NF-KAPPA B signaling pathway component expression FC differences between KC and LSCs.

**Supplementary Fig 3.**
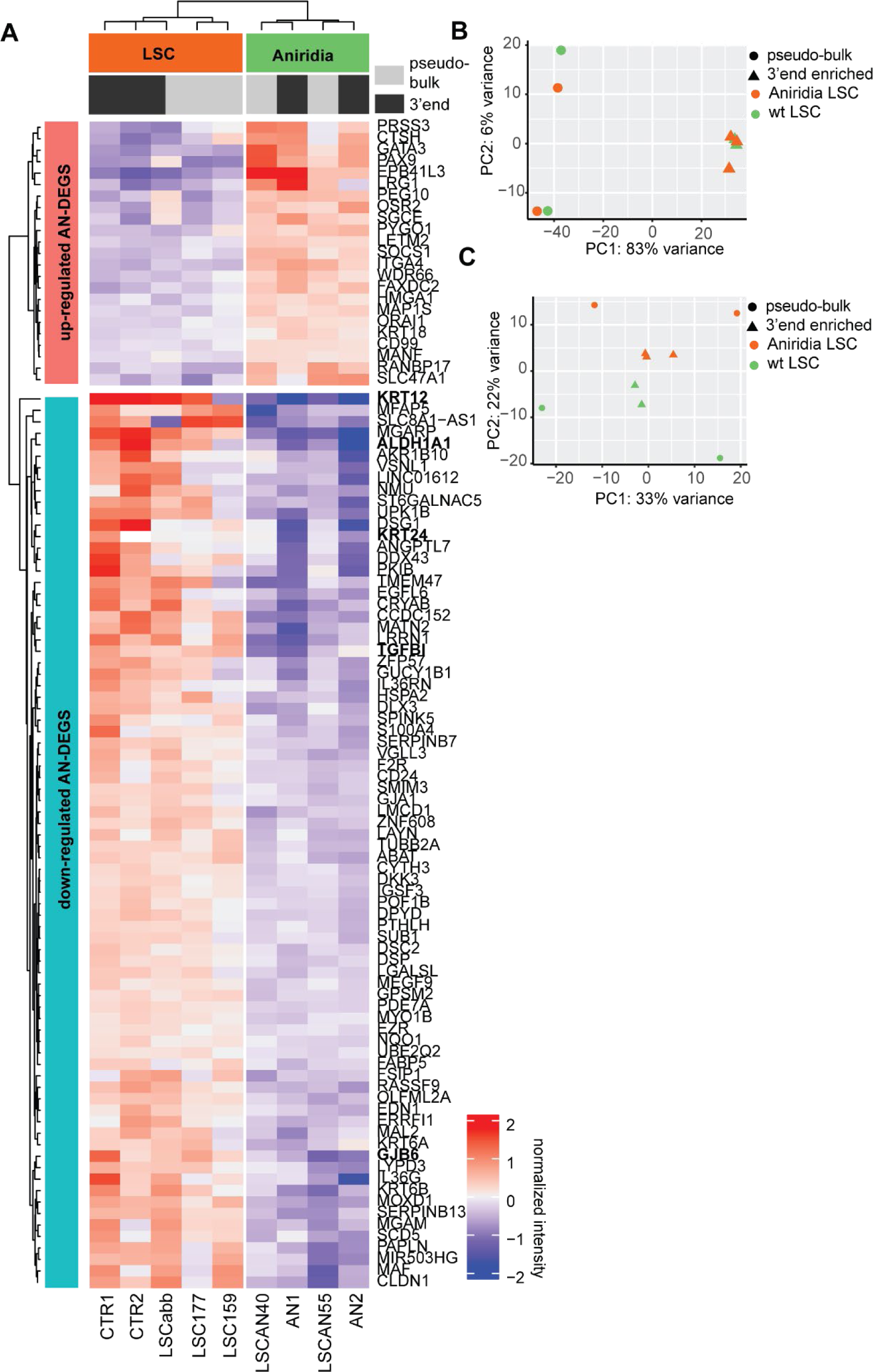
RNA-seq analysis of LSCs and Aniridia LSCs. (A) Heatmap of normalized DEG expression between control and aniridia patient LSCs (adjusted pval < 0.05), using k-means clustering with 2 clusters. (B) PCA plot of RNA-seq samples before batch correction. (C) PCA plot after batch correction.

**Supplementary Fig 4.**
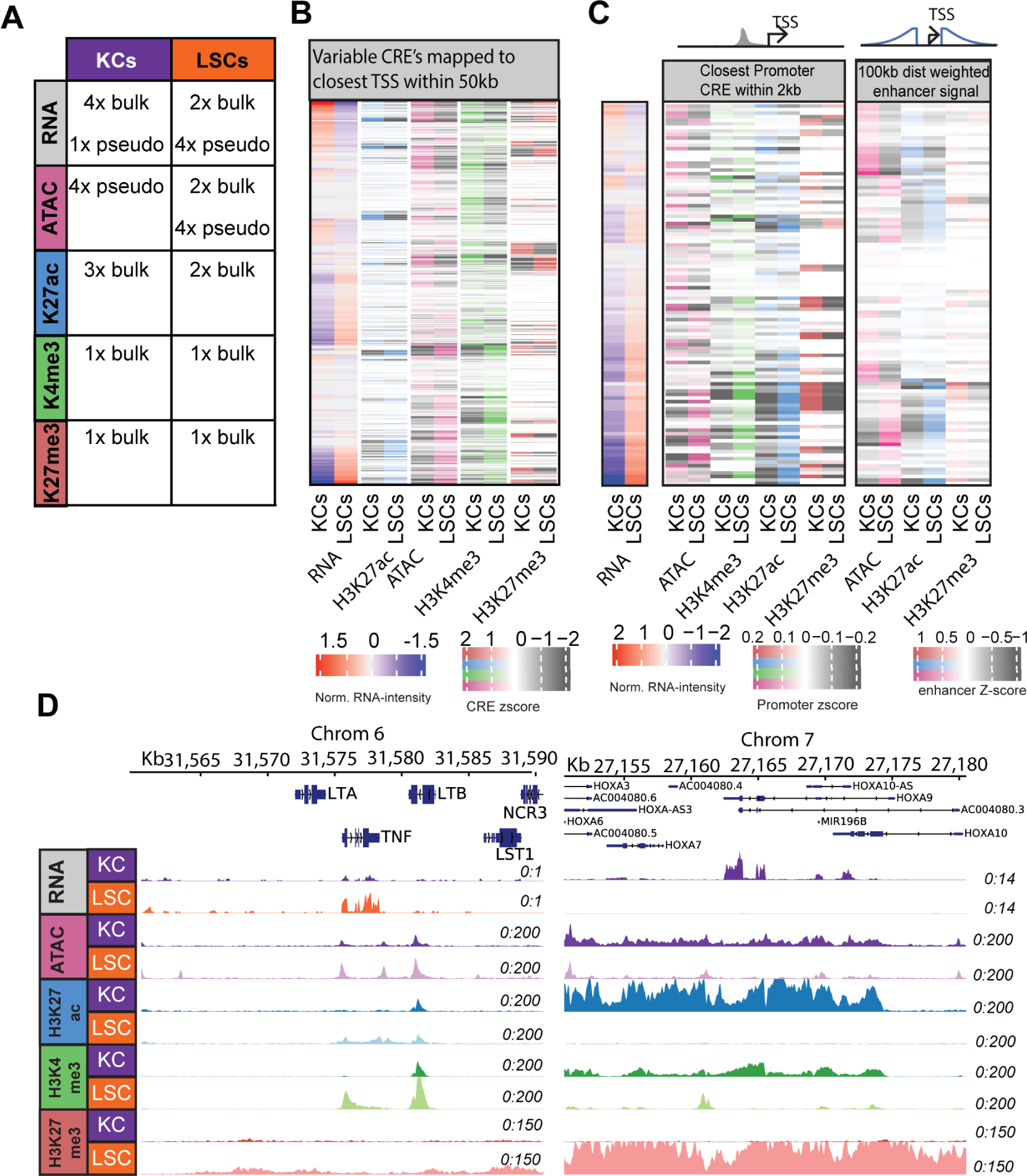
Additional vizualizations Regulatory Element (CRE) analysis. (A) Overview of data types used in our analysis. (B) Variable CREs mapped to the closest TSS within 50kb. Zscore normalized CRE signal intensities and normalized RNA-seq intensities. (C) Heatmap of PROGENy TNF and NF-KB target genes and the Z-score of the quantile normalized histone intensity signal of the closest CRE and the distance weighted enhancer signal. (D) TNF and HOXA9 TSS loci with signals of RNA-seq, ATAC-seq, ChIP-seq of H3K27ac, H3K4me3, and H3K27me3 in KCs and LSCs

**Supplementary Fig 5.**
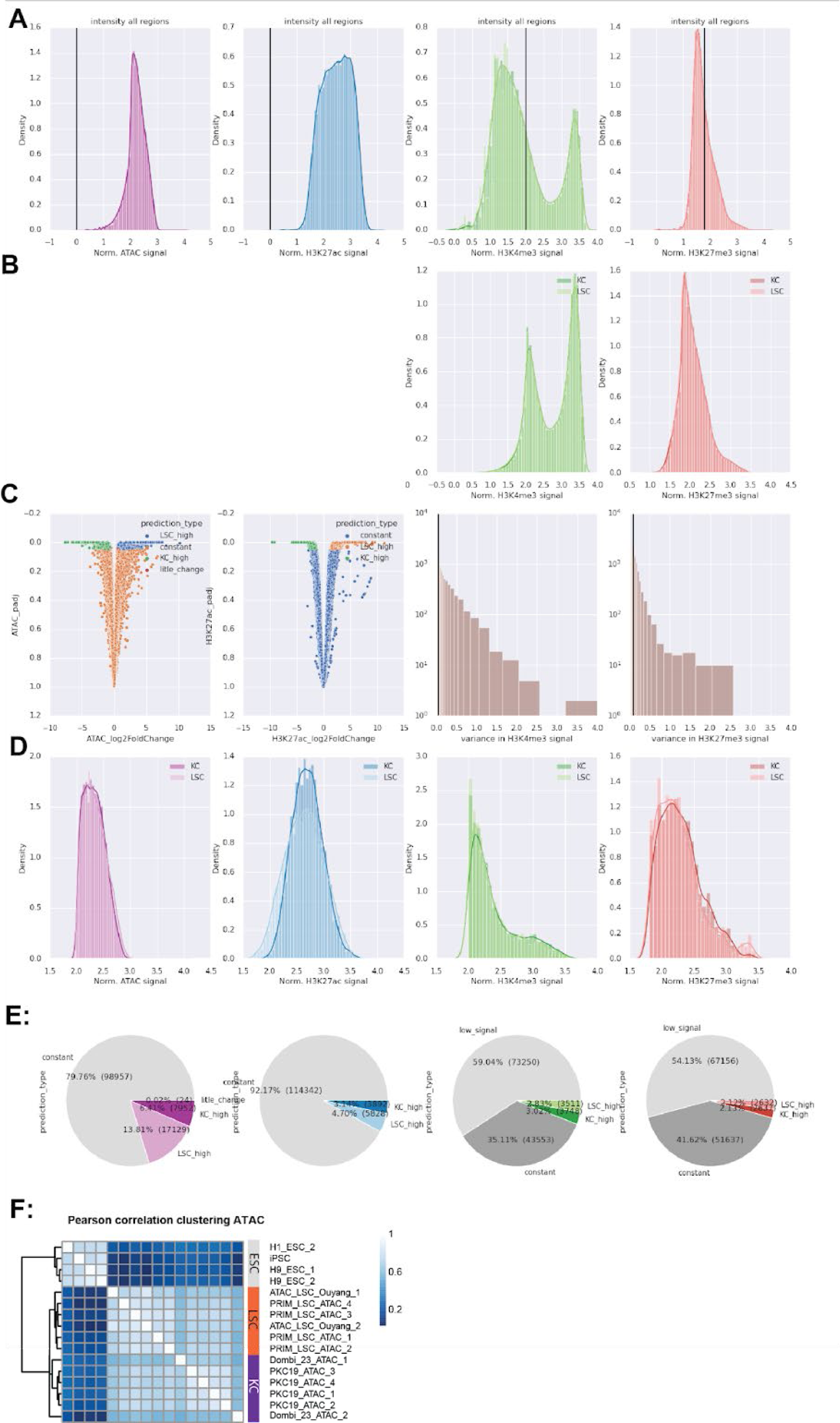
Additional quality control Regulatory Element (CRE) analysis. (A) quantile normalized intensity score of all ATAC peaks for the varying histone datasets. Including cutoff value for H3K4me3 and H3K27me3. (B) Resulting intensity score for H3K4me3 and H3k27me3 regions. (C) Deseq2 volcano plot of all ATAC & H3K27ac regions. Variance with the variance cutoff for H3K4me3 and H3K27me3. (D) resulting population of variable regions. E) Pie chart of region type distribution.

**Supplementary Fig 6.**
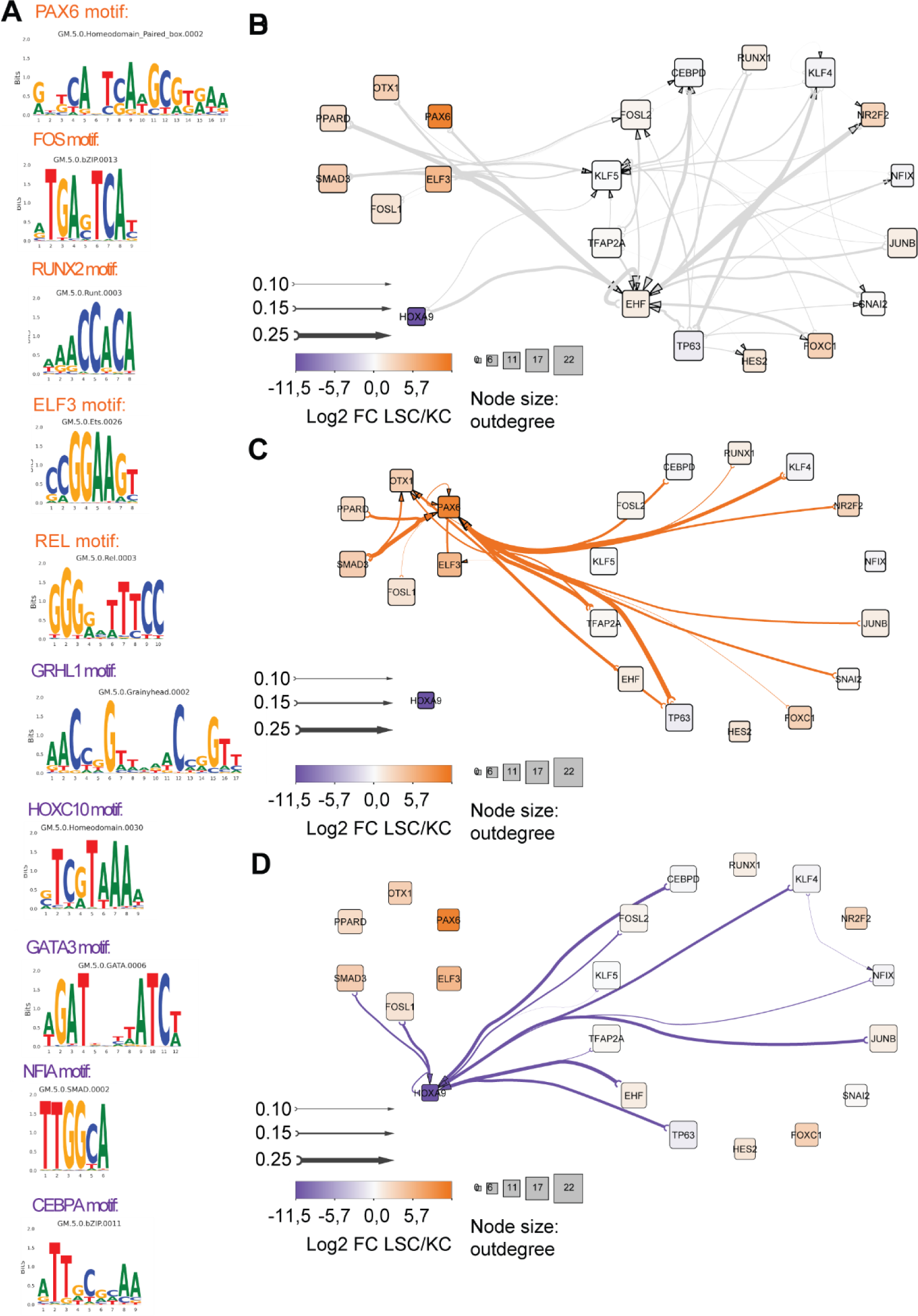
TFs Motifs and hierarchy vizualization. (A) Enriched motifs linked to the various TFs. (B) General epithelial interactions between TFs. Edge Width corresponds with Ananse binding score predictions. Node color represents RNAseq fold change between LSC and KCs, while node size represents outdegree. (C) Similar to B but with all the LSC specific interactions. (D) Similar to B but with all the KC specific interactions.

**Supplementary Fig 7.**
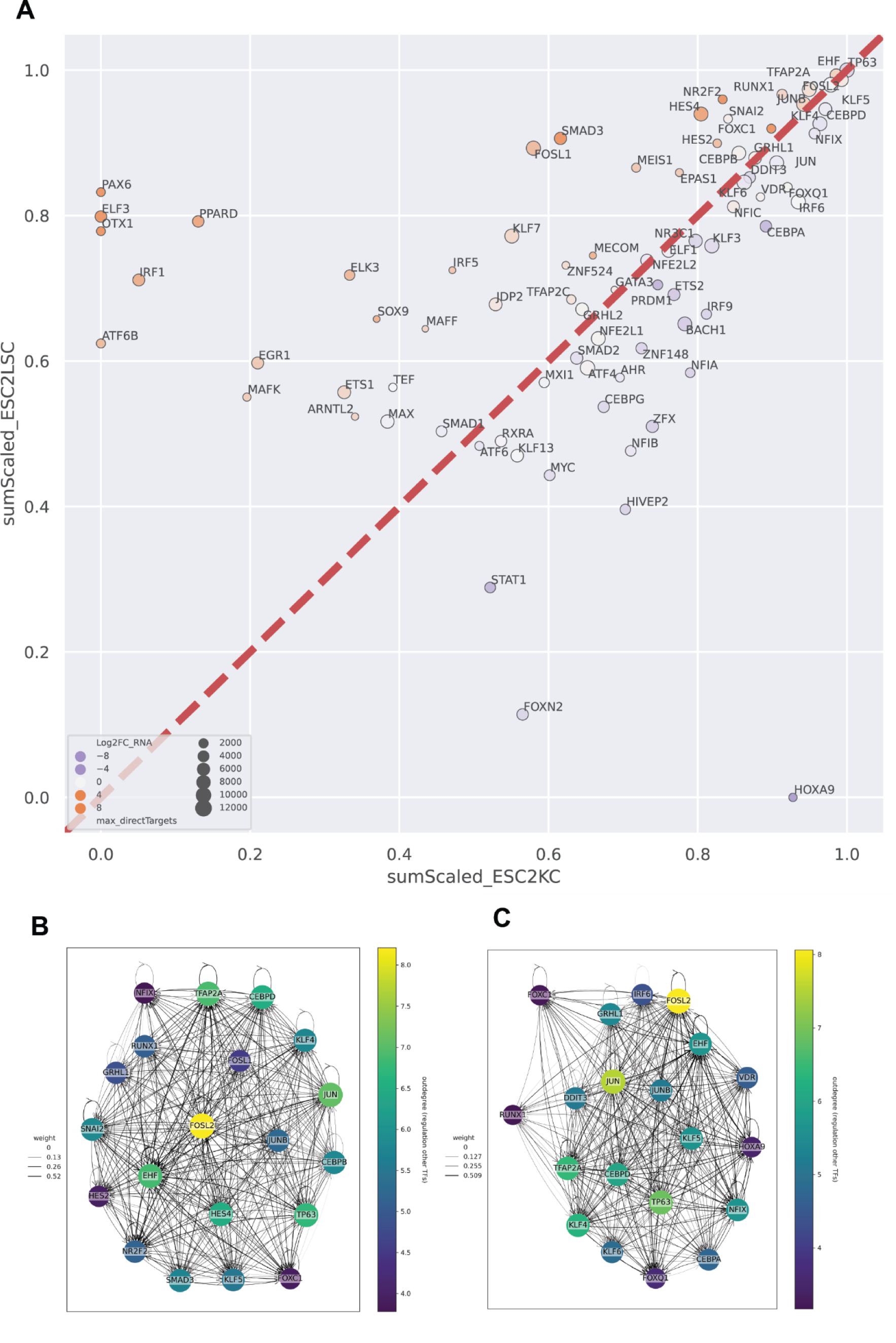
Extra labeled influence plot and outdegree analysis. (A) Ananse influence of ESC to KC (x-axis) and ESC to LSC (y-axis), circle size represents a maximum number of target genes in both comparisons. The circle color represents log2FC between LSC/KC. (B) ESC-LSC top TF interaction network generated by ANANSE. (C) ESC-KC top TF interaction network generated by ANANSE.

**Supplementary Fig 8.**
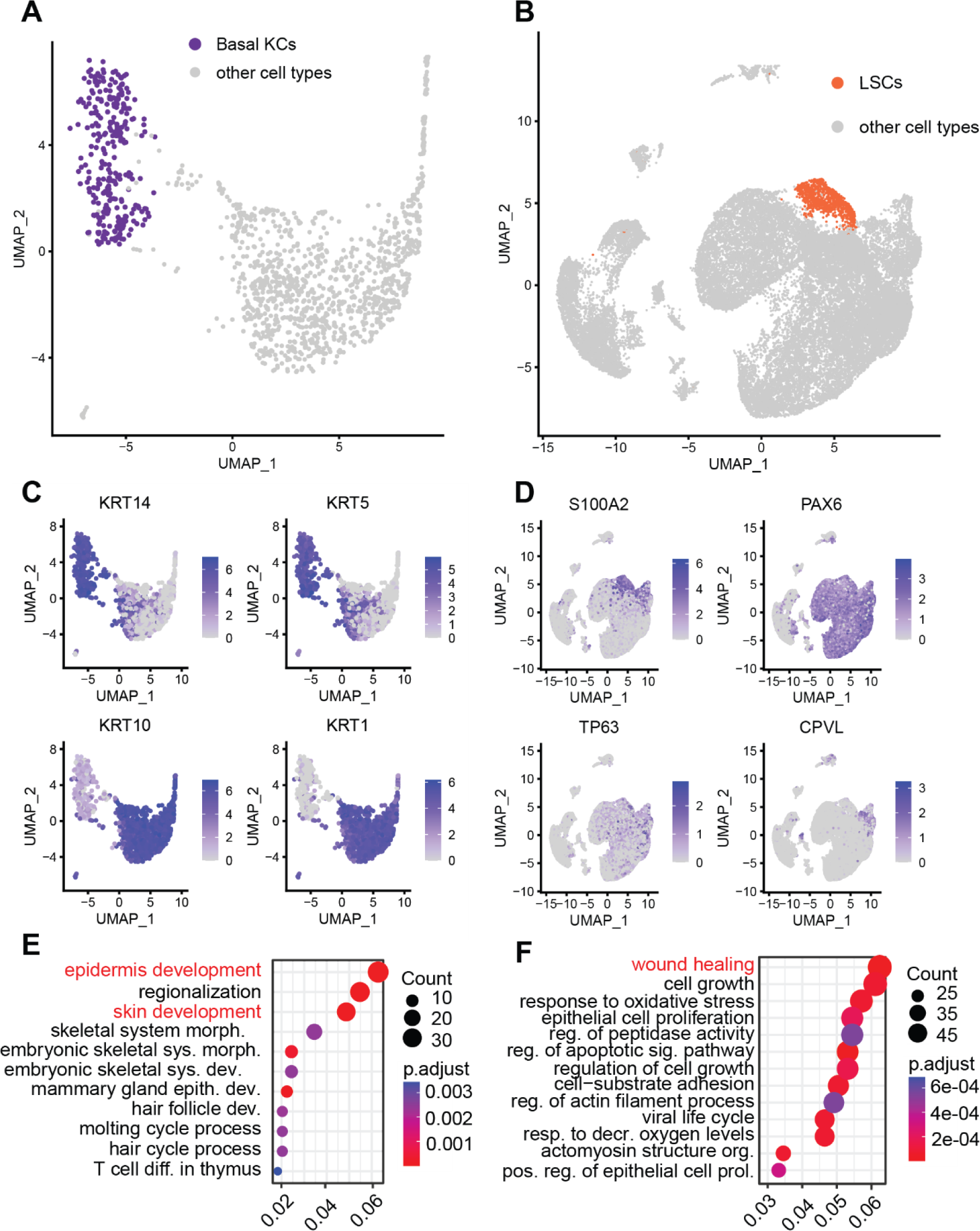
Additional quality control in vivo single-cell RNA-seq. (A) Umap of epidermal scRNAseq dataset of Atwood et al. Basal KC cluster used for validation is highlighted. (B) Umap of scCornea atlas of Collin et al, basal LSC cluster used for validation is highlighted. (C) Marker gene expression used to select the basal KCs cluster. (D) Marker gene expression used to select the basal LSC cluster. (E) GO-term enrichment of the basal-KC high DEGS enriched vs the human genome as a background and simplified using simplify. (F) GO-term enrichment of the basal-LSCs high DEGS enriched vs the human genome as a background and simplified using simplify.

**Supplementary Fig 9.**
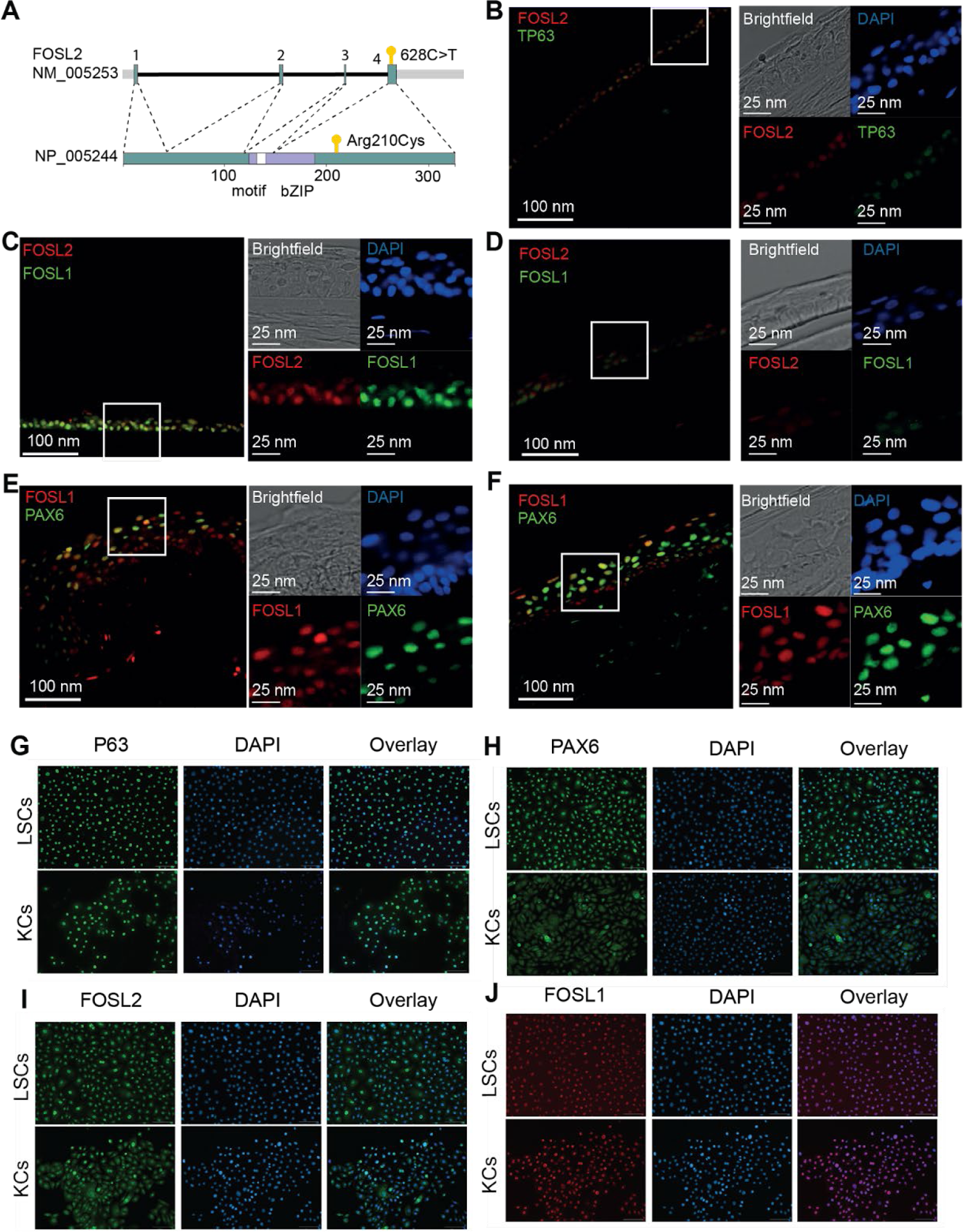
FOSL2 variant and protein stainings. (A) Overview of the FOSL2 transcript and protein, with the location of the variant of unknown significance. (B) FOSL2 and TP63 staining of the central cornea. (C) FOSL1 and FOSL2 staining of the peripheral cornea (D) FOSL1 and FOSL2 staining of the central cornea. (E) FOSL2 and PAX6 staining of the peripheral cornea. (F) FOSL1 and FOSL2 staining of the central cornea. (G-J) Immunocytochemistry analysis of fixed LSCs and KCs of transcription factors p63 (E), PAX6 (F), FOSL2 (G), and FOSL1 (H). Predicted general TFs p63, FOSL1 and FOSL2 are detected in the nuclei of both cell types while PAX6 is only detected in LSCs. Note that some non-nuclear signal is found in KCs stained with PAX6 but that is considered autofluorescence. DAPI staining (blue) depicts cell nuclei. Scale bar 100μm.

**Supplementary Fig 10.**
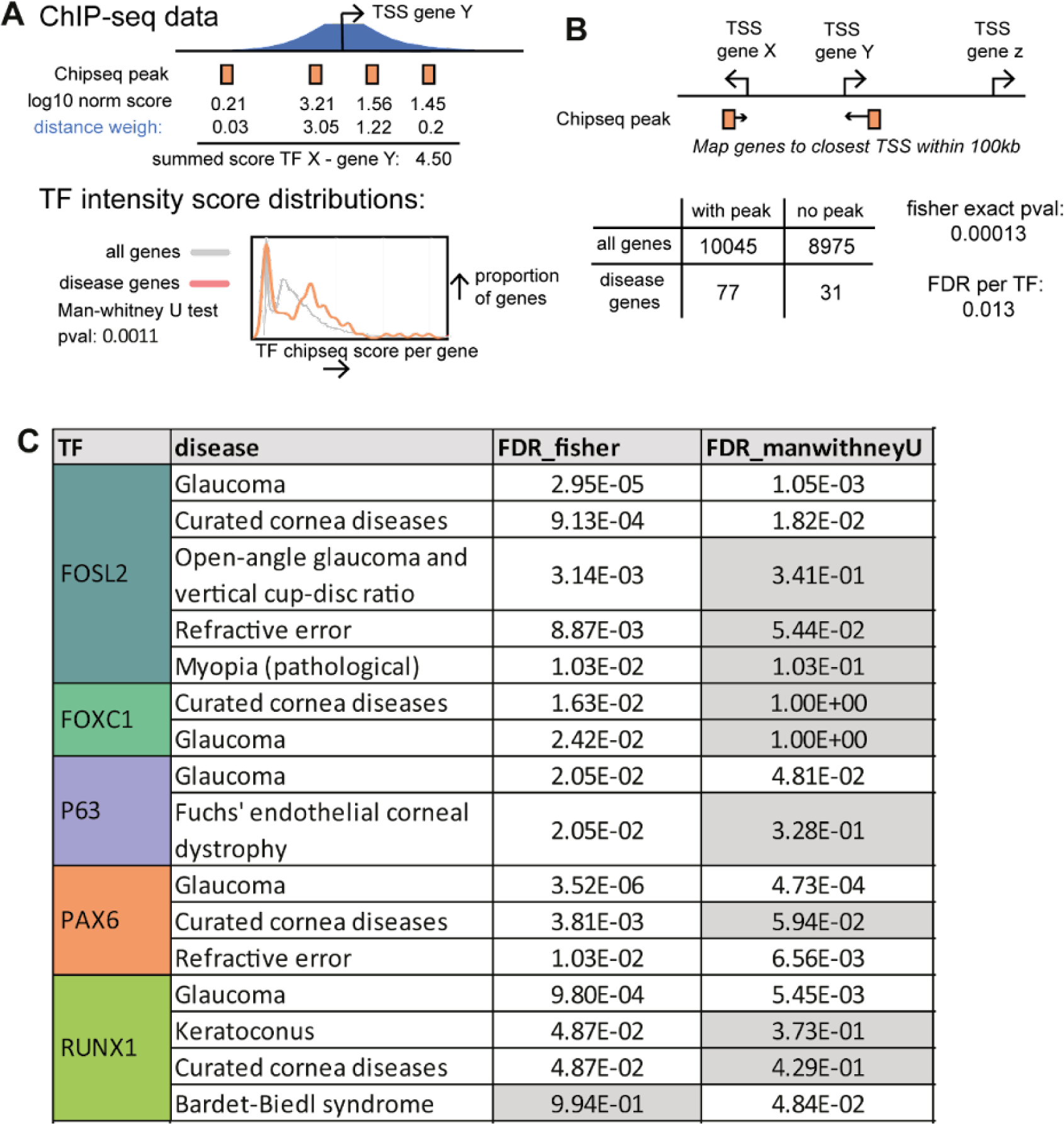
TF binding disease genes enrichment overview. (A) Approach for distance weighing and merging of TF ChIP-seqs per TF. This resulted in a TF-disease gene score distribution that was compared to the distribution of all genes with a one-sided Mann-Whitney U test. (B) Approach for linking the ChIP-seq peaks to the closest gene TSS, after which enrichment for disease genes was tested with a Fisher exact test. (C) FDR values of the significant enriched TFs resulting from the ChIP-seq Mann-Whitney U testsand the the Fisher exact test. Significant for FDR < 0.1.

**Supplementary Fig 11.**
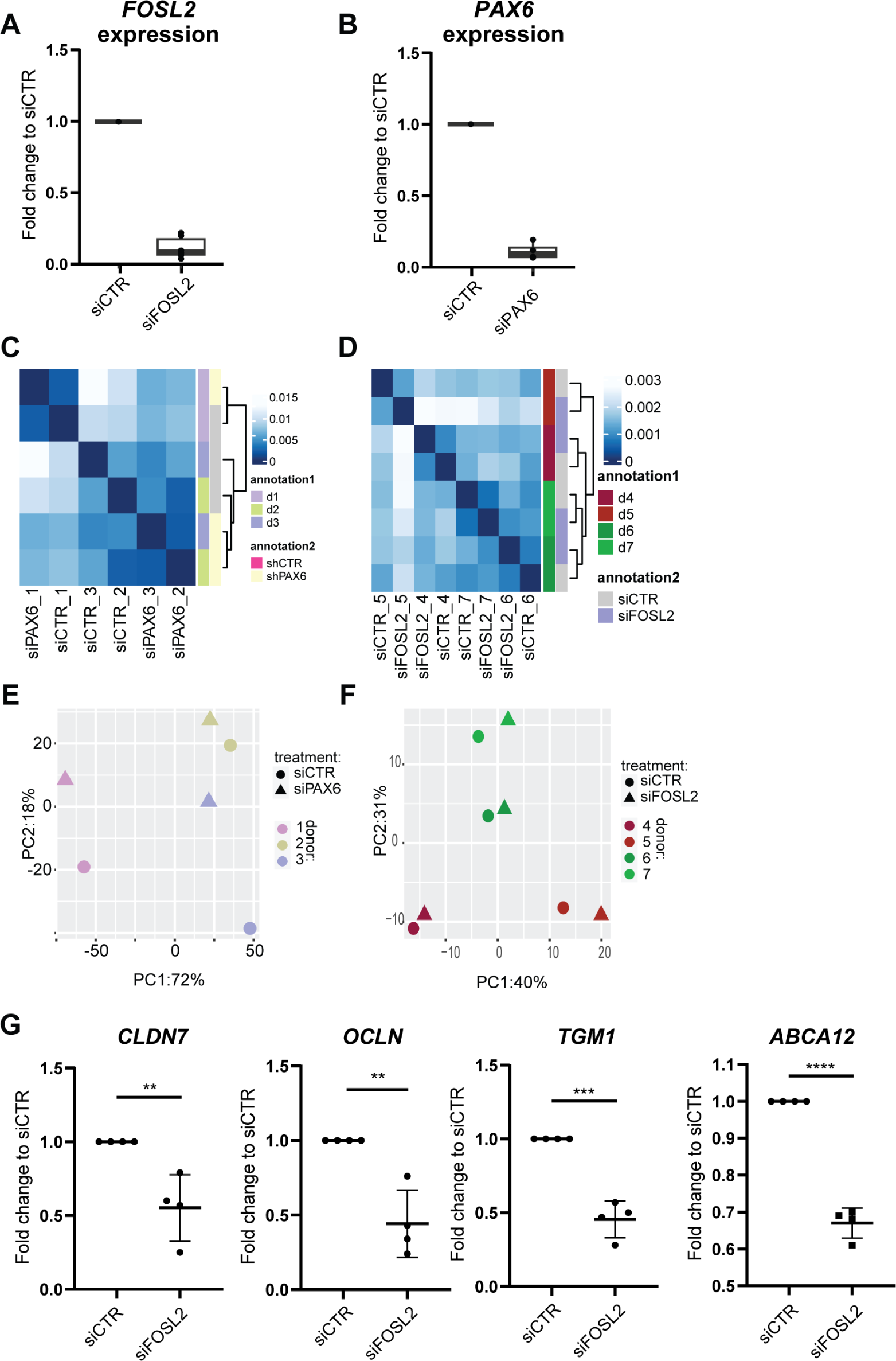
Additional quality control siRNA treatment and RNA sequencing. (A) qPCR validation FOSL2 knockdown. (B) qPCR validation PAX6 knockdown. (C) Pearson correlation matrix siCTR and siPAX6 samples. (D) Pearson correlation matrix siCTR and siFOSL2 samples. (E) PCA plot of RNAseq siPAX6 samples. (F) PCA plot of RNAseq siFOSL2 samples. (G) Transcripts CLDN7, OCLN, TGM1 and ABCA12 were measured in control LSCs (CTRL) and FOSL2 siRNA-knock down (FOSL2 KD) samples (n=4). Values represent fold change difference of FOSL2 KD to their respective CTRL and were normalized to internal housekeepers GAPDH and ACTB. (* pval<0.05, ** pval<0.01, *** pval<0.001, unpaired t-test analysis).

## References

1. Takahashi K, Yamanaka S. Induction of pluripotent stem cells from mouse embryonic and adult fibroblast cultures by defined factors. Cell. 25 augustus 2006;126(4):663-76.

2. Chambers SM, Studer L. Cell fate plug and play: direct reprogramming and induced pluripotency. Cell. 10 juni 2011;145(6):827-30.

3. Li M, Belmonte JCI. Ground rules of the pluripotency gene regulatory network. Nat Rev Genet. maart 2017;18(3):180–91.

4. Zaret KS. Pioneer Transcription Factors Initiating Gene Network Changes. Annu Rev Genet. 23 november 2020;54:367–85.

5. Epstein DJ. Cis-regulatory mutations in human disease. Brief Funct Genomic Proteomic. juli 2009;8(4):310–6.

6. Lee TI, Young RA. Transcriptional Regulation and its Misregulation in Disease. Cell. 14 maart 2013;152(6):1237-51.

7. Roberts N, Horsley V. Developing stratified epithelia: lessons from the epidermis and thymus. Wiley Interdiscip Rev Dev Biol. december 2014;3(6):389–402.

8. Donati G, Watt FM. Stem cell heterogeneity and plasticity in epithelia. Cell Stem Cell. 7 mei 2015;16(5):465-76.

9. Bashir H, Seykora JT, Lee V. Invisible Shield: Review of the Corneal Epithelium as a Barrier to UV Radiation, Pathogens, and Other Environmental Stimuli. J Ophthalmic Vis Res. 2017;12(3):305–11.

10. Gonzalez G, Sasamoto Y, Ksander BR, Frank MH, Frank NY. Limbal Stem Cells: Identity, Developmental Origin and Therapeutic Potential. Wiley Interdiscip Rev Dev Biol [Internet]. maart 2018 [geciteerd 21 april 2021];7(2). Beschikbaar op: https://www.ncbi.nlm.nih.gov/pmc/articles/PMC5814333/

11. Bhaduri A, Ungewickell A, Boxer LD, Lopez-Pajares V, Zarnegar BJ, Khavari PA. Network Analysis Identifies Mitochondrial Regulation of Epidermal Differentiation by MPZL3 and FDXR. Dev Cell. 23 november 2015;35(4):444–57.

12. Rubin AJ, Barajas BC, Furlan-Magaril M, Lopez-Pajares V, Mumbach MR, Howard I, e.a. Lineage-specific dynamic and pre-established enhancer–promoter contacts cooperate in terminal differentiation. Nat Genet. oktober 2017;49(10):1522–8.

13. Soares E, Zhou H. Master regulatory role of p63 in epidermal development and disease. Cell Mol Life Sci CMLS. april 2018;75(7):1179–90.

14. Li L, Wang Y, Torkelson JL, Shankar G, Pattison JM, Zhen HH, e.a. TFAP2C- and p63-Dependent Networks Sequentially Rearrange Chromatin Landscapes to Drive Human Epidermal Lineage Commitment. Cell Stem Cell. 7 februari 2019;24(2):271-284.e8.

15. Sen GL, Boxer LD, Webster DE, Bussat RT, Qu K, Zarnegar BJ, e.a. ZNF750 is a p63 target gene that induces KLF4 to drive terminal epidermal differentiation. Dev Cell. 13 maart 2012;22(3):669-77.

16. Qu J, Tanis SEJ, Smits JPH, Kouwenhoven EN, Oti M, van den Bogaard EH, e.a. Mutant p63 Affects Epidermal Cell Identity through Rewiring the Enhancer Landscape. Cell Rep. 18 december 2018;25(12):3490–3503.e4.

17. Kouwenhoven, E.N., Oti, M., Niehues, H., Heeringen, S.J. van, Schalkwijk, J., Stunnenberg, H.G., e.a. Transcription factor p63 bookmarks and regulates dynamic enhancers during epidermal differentiation. EMBO Rep. 2015;16:863–78.

18. Qu J, Yi G, Zhou H. p63 cooperates with CTCF to modulate chromatin architecture in skin keratinocytes. Epigenetics Chromatin. 4 juni 2019;12(1):31.

19. van Bokhoven H, Brunner HG. Splitting p63. Am J Hum Genet. juli 2002;71(1):1–13.

20. Rinne T, Hamel B, van Bokhoven H, Brunner HG. Pattern of p63 mutations and their phenotypes--update. Am J Med Genet A. 1 juli 2006;140(13):1396-406.

21. Rinne T, Brunner HG, van Bokhoven H. p63-associated disorders. Cell Cycle Georget Tex. 1 februari 2007;6(3):262-8.

22. Hanson I, Van Heyningen V. Pax6: more than meets the eye. Trends Genet TIG. juli 1995;11(7):268–72.

23. Ashery-Padan R, Gruss P. Pax6 lights-up the way for eye development. Curr Opin Cell Biol. december 2001;13(6):706–14.

24. Cvekl A, Callaerts P. PAX6: 25th anniversary and more to learn. Exp Eye Res. maart 2017;156:10–21.

25. Xie Q, Cvekl A. The orchestration of mammalian tissue morphogenesis through a series of coherent feed-forward loops. J Biol Chem. 16 december 2011;286(50):43259–71.

26. Cohen-Tayar Y, Cohen H, Mitiagin Y, Abravanel Z, Levy C, Idelson M, e.a. Pax6 regulation of Sox9 in the mouse retinal pigmented epithelium controls its timely differentiation and choroid vasculature development. Dev Camb Engl. 9 augustus 2018;145(15):dev163691.

27. Ypsilanti AR, Pattabiraman K, Catta-Preta R, Golonzhka O, Lindtner S, Tang K, e.a. Transcriptional network orchestrating regional patterning of cortical progenitors. Proc Natl Acad Sci U S A. 21 december 2021;118(51):e2024795118.

28. van Heyningen V, Williamson KA. PAX6 in sensory development. Hum Mol Genet. 15 mei 2002;11(10):1161-7.

29. Shaham O, Menuchin Y, Farhy C, Ashery-Padan R. Pax6: a multi-level regulator of ocular development. Prog Retin Eye Res. september 2012;31(5):351–76.

30. Ouyang H, Xue Y, Lin Y, Zhang X, Xi L, Patel S, e.a. WNT7A and PAX6 define corneal epithelium homeostasis and pathogenesis. Nature. 17 juli 2014;511(7509):358-61.

31. Lagali N, Wowra B, Fries FN, Latta L, Moslemani K, Utheim TP, e.a. Early phenotypic features of aniridia-associated keratopathy and association with PAX6 coding mutations. Ocul Surf. januari 2020;18(1):130–40.

32. Li M, Huang H, Li L, He C, Zhu L, Guo H, e.a. Core transcription regulatory circuitry orchestrates corneal epithelial homeostasis. Nat Commun. 18 januari 2021;12(1):420.

33. Lima Cunha D, Arno G, Corton M, Moosajee M. The Spectrum of PAX6 Mutations and Genotype-Phenotype Correlations in the Eye. Genes [Internet]. 17 december 2019 [geciteerd 3 maart 2021];10(12). Beschikbaar op: https://www.ncbi.nlm.nih.gov/pmc/articles/PMC6947179/

34. Kit V, Cunha DL, Hagag AM, Moosajee M. Longitudinal genotype-phenotype analysis in 86 patients with PAX6-related aniridia. JCI Insight. 22 juli 2021;6(14):148406.

35. Rinne T, Clements SE, Lamme E, Duijf PHG, Bolat E, Meijer R, e.a. A novel translation re-initiation mechanism for the p63 gene revealed by amino-terminal truncating mutations in Rapp-Hodgkin/Hay-Wells-like syndromes. Hum Mol Genet. 1 juli 2008;17(13):1968-77.

36. Di Iorio E, Kaye SB, Ponzin D, Barbaro V, Ferrari S, Böhm E, e.a. Limbal stem cell deficiency and ocular phenotype in ectrodactyly-ectodermal dysplasia-clefting syndrome caused by p63 mutations. Ophthalmology. januari 2012;119(1):74–83.

37. Tümer Z, Bach-Holm D. Axenfeld–Rieger syndrome and spectrum of PITX2 and FOXC1 mutations. Eur J Hum Genet. december 2009;17(12):1527–39.

38. Seo S, Singh HP, Lacal PM, Sasman A, Fatima A, Liu T, e.a. Forkhead box transcription factor FoxC1 preserves corneal transparency by regulating vascular growth. Proc Natl Acad Sci. 7 februari 2012;109(6):2015-20.

39. Li M, Zhu L, Liu J, Huang H, Guo H, Wang L, e.a. Loss of FOXC1 contributes to the corneal epithelial fate switch and pathogenesis. Signal Transduct Target Ther. 8 januari 2021;6(1):1-11.

40. Kitazawa K, Hikichi T, Nakamura T, Sotozono C, Kinoshita S, Masui S. PAX6 regulates human corneal epithelium cell identity. Exp Eye Res. 1 januari 2017;154:30-8.

41. van den Bogaard EH, Rodijk-Olthuis D, Jansen PAM, van Vlijmen-Willems IMJJ, van Erp PE, Joosten I, e.a. Rho kinase inhibitor Y-27632 prolongs the life span of adult human keratinocytes, enhances skin equivalent development, and facilitates lentiviral transduction. Tissue Eng Part A. september 2012;18(17-18):1827–36.

42. Lužnik Z, Bertolin M, Breda C, Ferrari B, Barbaro V, Schollmayer P, e.a. Preservation of Ocular Epithelial Limbal Stem Cells: The New Frontier in Regenerative Medicine. Adv Exp Med Biol. 2016;951:179–89.

43. Gene Ontology Consortium. The Gene Ontology (GO) database and informatics resource. Nucleic Acids Res. 1 januari 2004;32(suppl_1):D258-61.

44. Wu T, Hu E, Xu S, Chen M, Guo P, Dai Z, e.a. clusterProfiler 4.0: A universal enrichment tool for interpreting omics data. Innov N Y N. 28 augustus 2021;2(3):100141.

45. Subramanian A, Tamayo P, Mootha VK, Mukherjee S, Ebert BL, Gillette MA, e.a. Gene set enrichment analysis: A knowledge-based approach for interpreting genome-wide expression profiles. Proc Natl Acad Sci. 25 oktober 2005;102(43):15545-50.

46. Liberzon A, Birger C, Thorvaldsdóttir H, Ghandi M, Mesirov JP, Tamayo P. The Molecular Signatures Database (MSigDB) hallmark gene set collection. Cell Syst. 23 december 2015;1(6):417–25.

47. Schubert M, Klinger B, Klünemann M, Sieber A, Uhlitz F, Sauer S, e.a. Perturbation-response genes reveal signaling footprints in cancer gene expression. Nat Commun. 2 januari 2018;9(1):20.

48. Kanehisa M, Goto S. KEGG: Kyoto Encyclopedia of Genes and Genomes. Nucleic Acids Res. 1 januari 2000;28(1):27-30.

49. Latta L, Nordström K, Stachon T, Langenbucher A, Fries FN, Szentmáry N, e.a. Expression of retinoic acid signaling components ADH7 and ALDH1A1 is reduced in aniridia limbal epithelial cells and a siRNA primary cell based aniridia model. Exp Eye Res. februari 2019;179:8–17.

50. Roux LN, Petit I, Domart R, Concordet JP, Qu J, Zhou H, e.a. Modeling of Aniridia-Related Keratopathy by CRISPR/Cas9 Genome Editing of Human Limbal Epithelial Cells and Rescue by Recombinant PAX6 Protein. STEM CELLS. 2018;36(9):1421–9.

51. Bruse N, Heeringen SJ van. GimmeMotifs: an analysis framework for transcription factor motif analysis [Internet]. bioRxiv; 2018 [geciteerd 21 december 2022]. p. 474403. Beschikbaar op: https://www.biorxiv.org/content/10.1101/474403v1

52. Xu Q, Georgiou G, Frölich S, van der Sande M, Veenstra GJC, Zhou H, e.a. ANANSE: an enhancer network-based computational approach for predicting key transcription factors in cell fate determination. Nucleic Acids Res. 20 augustus 2021;49(14):7966-85.

53. Hawkins RD, Hon GC, Lee LK, Ngo Q, Lister R, Pelizzola M, e.a. Distinct epigenomic landscapes of pluripotent and lineage-committed human cells. Cell Stem Cell. 7 mei 2010;6(5):479-91.

54. Wang S, Drummond ML, Guerrero-Juarez CF, Tarapore E, MacLean AL, Stabell AR, e.a. Single cell transcriptomics of human epidermis identifies basal stem cell transition states. Nat Commun. 25 augustus 2020;11(1):4239.

55. Collin J, Queen R, Zerti D, Bojic S, Dorgau B, Moyse N, e.a. A single cell atlas of human cornea that defines its development, limbal progenitor cells and their interactions with the immune cells. Ocul Surf. juli 2021;21:279–98.

56. Yuan J, Chen F, Fan D, Jiang Q, Xue Z, Zhang J, e.a. EyeDiseases: an integrated resource for dedicating to genetic variants, gene expression and epigenetic factors of human eye diseases. NAR Genomics Bioinforma. 1 juni 2021;3(2):lqab050.

57. Klein RH, Lin Z, Hopkin AS, Gordon W, Tsoi LC, Liang Y, e.a. GRHL3 binding and enhancers rearrange as epidermal keratinocytes transition between functional states. PLOS Genet. 26 april 2017;13(4):e1006745.

58. Gong D, Yan C, Yu F, Yan D, Wu N, Chen L, e.a. Direct oral mucosal epithelial transplantation supplies stem cells and promotes corneal wound healing to treat refractory persistent corneal epithelial defects. Exp Eye Res. februari 2022;215:108934.

59. Hassan NT, AbdelAziz NA. Oral Mucosal Stem Cells, Human Immature Dental Pulp Stem Cells and Hair Follicle Bulge Stem Cells as Adult Stem Cells Able to Correct Limbal Stem Cell Deficiency. Curr Stem Cell Res Ther. 2018;13(5):356–61.

60. Simeone A. Otx1 and Otx2 in the development and evolution of the mammalian brain. EMBO J. 1 december 1998;17(23):6790–8.

61. Huang B, Li X, Tu X, Zhao W, Zhu D, Feng Y, e.a. OTX1 regulates cell cycle progression of neural progenitors in the developing cerebral cortex. J Biol Chem. 9 februari 2018;293(6):2137-48.

62. Martinez-Morales JR, Signore M, Acampora D, Simeone A, Bovolenta P. Otx genes are required for tissue specification in the developing eye. Development. 1 juni 2001;128(11):2019-30.

63. Samuel A, Housset M, Fant B, Lamonerie T. Otx2 ChIP-seq Reveals Unique and Redundant Functions in the Mature Mouse Retina. PLOS ONE. 18 februari 2014;9(2):e89110.

64. Yoshida N, Yoshida S, Araie M, Handa H, Nabeshima Y ichi. Ets family transcription factor ESE-1 is expressed in corneal epithelial cells and is involved in their differentiation. Mech Dev. 1 oktober 2000;97(1):27-34.

65. Karin M, Liu Z g, Zandi E. AP-1 function and regulation. Curr Opin Cell Biol. april 1997;9(2):240–6.

66. Eckert RL, Adhikary G, Young CA, Jans R, Crish JF, Xu W, e.a. AP1 transcription factors in epidermal differentiation and skin cancer. J Skin Cancer. 2013;2013:537028.

67. Borrelli S, Testoni B, Callari M, Alotto D, Castagnoli C, Romano RA, e.a. Reciprocal regulation of p63 by C/EBP delta in human keratinocytes. BMC Mol Biol. 28 september 2007;8(1):85.

68. Candi E, Terrinoni A, Rufini A, Chikh A, Lena AM, Suzuki Y, e.a. p63 is upstream of IKK alpha in epidermal development. J Cell Sci. 15 november 2006;119(Pt 22):4617–22.

69. Chikh A, Sayan E, Thibaut S, Lena AM, DiGiorgi S, Bernard BA, e.a. Expression of GATA-3 in epidermis and hair follicle: relationship to p63. Biochem Biophys Res Commun. 14 september 2007;361(1):1–6.

70. Zeitvogel J, Jokmin N, Rieker S, Klug I, Brandenberger C, Werfel T. GATA3 regulates FLG and FLG2 expression in human primary keratinocytes. Sci Rep. 19 september 2017;7(1):11847.

71. Rinn JL, Wang JK, Allen N, Brugmann SA, Mikels AJ, Liu H, e.a. A dermal HOX transcriptional program regulates site-specific epidermal fate. Genes Dev. 1 februari 2008;22(3):303-7.

72. Gehring WJ. The animal body plan, the prototypic body segment, and eye evolution. Evol Dev. 2012;14(1):34–46.

73. Kitazawa K, Hikichi T, Nakamura T, Mitsunaga K, Tanaka A, Nakamura M, e.a. OVOL2 Maintains the Transcriptional Program of Human Corneal Epithelium by Suppressing Epithelial-to-Mesenchymal Transition. Cell Rep. 10 mei 2016;15(6):1359-68.

74. Menzel-Severing J, Zenkel M, Polisetti N, Sock E, Wegner M, Kruse FE, e.a. Transcription factor profiling identifies Sox9 as regulator of proliferation and differentiation in corneal epithelial stem/progenitor cells. Sci Rep. 6 juli 2018;8(1):10268.

75. McConnell BB, Ghaleb AM, Nandan MO, Yang VW. The diverse functions of Krüppel-like factors 4 and 5 in epithelial biology and pathobiology. BioEssays News Rev Mol Cell Dev Biol. juni 2007;29(6):549–57.

76. Kenchegowda D, Harvey SAK, Swamynathan S, Lathrop KL, Swamynathan SK. Critical Role of Klf5 in Regulating Gene Expression during Post-Eyelid Opening Maturation of Mouse Corneas. PLoS ONE. 14 september 2012;7(9):e44771.

77. Stephens DN, Klein RH, Salmans ML, Gordon W, Ho H, Andersen B. The Ets transcription factor EHF as a regulator of cornea epithelial cell identity. J Biol Chem. 2013/10/18 dr. 29 november 2013;288(48):34304-24.

78. Tiwari A, Loughner CL, Swamynathan S, Swamynathan SK. KLF4 Plays an Essential Role in Corneal Epithelial Homeostasis by Promoting Epithelial Cell Fate and Suppressing Epithelial– Mesenchymal Transition. Invest Ophthalmol Vis Sci. mei 2017;58(5):2785–95.

79. Cieślar-Pobuda A, Rafat M, Knoflach V, Skonieczna M, Hudecki A, Małecki A, e.a. Human induced pluripotent stem cell differentiation and direct transdifferentiation into corneal epithelial-like cells. Oncotarget. 2 juni 2016;7(27):42314-29.

80. Wang X, Shan X, Gregory-Evans CY. A mouse model of aniridia reveals the in vivo downstream targets of Pax6 driving iris and ciliary body development in the eye. Biochim Biophys Acta Mol Basis Dis. januari 2017;1863(1):60–7.

81. Shetty A, Tripathi SK, Junttila S, Buchacher T, Biradar R, Bhosale SD, e.a. A systematic comparison of FOSL1, FOSL2 and BATF-mediated transcriptional regulation during early human Th17 differentiation. Nucleic Acids Res. 20 mei 2022;50(9):4938-58.

82. Shinde V, Hu N, Mahale A, Maiti G, Daoud Y, Eberhart CG, e.a. RNA sequencing of corneas from two keratoconus patient groups identifies potential biomarkers and decreased NRF2-antioxidant responses. Sci Rep. 18 juni 2020;10(1):9907.

83. McHenry JZ, Leon A, Matthaei KI, Cohen DR. Overexpression of fra-2 in transgenic mice perturbs normal eye development. Oncogene. september 1998;17(9):1131–40.

84. Ferreira MAR, Vonk JM, Baurecht H, Marenholz I, Tian C, Hoffman JD, e.a. Eleven loci with new reproducible genetic associations with allergic disease risk. J Allergy Clin Immunol. 1 februari 2019;143(2):691-9.

85. Cospain A, Rivera-Barahona A, Dumontet E, Gener B, Bailleul-Forestier I, Meyts I, e.a. FOSL2 truncating variants in the last exon cause a neurodevelopmental disorder with scalp and enamel defects. Genet Med. 1 december 2022;24(12):2475–86.

86. Wan X, Guan S, Hou Y, Qin Y, Zeng H, Yang L, e.a. FOSL2 promotes VEGF-independent angiogenesis by transcriptionnally activating Wnt5a in breast cancer-associated fibroblasts. Theranostics. 5 maart 2021;11(10):4975-91.

87. Guo J, Shen S, Liu X, Ruan X, Zheng J, Liu Y, e.a. Role of linc00174/miR-138-5p (miR-150-5p)/FOSL2 Feedback Loop on Regulating the Blood-Tumor Barrier Permeability. Mol Ther - Nucleic Acids. 6 december 2019;18:1072–90.

88. Diverse Regulation of Claudin-1 and Claudin-4 in Atopic Dermatitis - ScienceDirect [Internet]. [geciteerd 21 juni 2023]. Beschikbaar op: https://www.sciencedirect.com/science/article/pii/S0002944015004277?via%3Dihub

89. Kirschner N, Poetzl C, von den Driesch P, Wladykowski E, Moll I, Behne MJ, e.a. Alteration of Tight Junction Proteins Is an Early Event in Psoriasis: Putative Involvement of Proinflammatory Cytokines. Am J Pathol. 1 september 2009;175(3):1095–106.

90. Latta L, Viestenz A, Stachon T, Colanesi S, Szentmáry N, Seitz B, e.a. Human aniridia limbal epithelial cells lack expression of keratins K3 and K12. Exp Eye Res. februari 2018;167:100–9.

91. Schlötzer-Schrehardt U, Latta L, Gießl A, Zenkel M, Fries FN, Käsmann-Kellner B, e.a. Dysfunction of the limbal epithelial stem cell niche in aniridia-associated keratopathy. Ocul Surf. 1 juli 2021;21:160-73.

92. Fasolo A, Pedrotti E, Passilongo M, Marchini G, Monterosso C, Zampini R, e.a. Safety outcomes and long-term effectiveness of ex vivo autologous cultured limbal epithelial transplantation for limbal stem cell deficiency. Br J Ophthalmol. mei 2017;101(5):640–9.

93. Lužnik Z, Breda C, Barbaro V, Ferrari S, Migliorati A, Di Iorio E, e.a. Towards xeno-free cultures of human limbal stem cells for ocular surface reconstruction. Cell Tissue Bank. december 2017;18(4):461–74.

94. Hashimshony T, Senderovich N, Avital G, Klochendler A, de Leeuw Y, Anavy L, e.a. CEL-Seq2: sensitive highly-multiplexed single-cell RNA-Seq. Genome Biol. 28 april 2016;17(1):77.

95. Kouwenhoven EN, Heeringen SJ van, Tena JJ, Oti M, Dutilh BE, Alonso ME, e.a. Genome-Wide Profiling of p63 DNA–Binding Sites Identifies an Element that Regulates Gene Expression during Limb Development in the 7q21 SHFM1 Locus. PLOS Genet. 19 augustus 2010;6(8):e1001065.

96. Lima Cunha D, Oram A, Gruber R, Plank R, Lingenhel A, Gupta MK, e.a. hiPSC-Derived Epidermal Keratinocytes from Ichthyosis Patients Show Altered Expression of Cornification Markers. Int J Mol Sci. 11 februari 2021;22(4):1785.

97. Maarten-vd-Sande, Siebren Frölich, Jos Smits, Simon van Heeringen, Quan Xu, mkolmus. vanheeringen-lab/seq2science: Release v0.1.0 [Internet]. Zenodo; 2020 [geciteerd 30 juli 2020]. Beschikbaar op: https://zenodo.org/record/3946493#.XyKceSgzYuU

98. Chen S, Zhou Y, Chen Y, Gu J. fastp: an ultra-fast all-in-one FASTQ preprocessor. Bioinformatics. 1 september 2018;34(17):i884–90.

99. Heeringen SJ van. genomepy: download genomes the easy way. J Open Source Softw. 15 augustus 2017;2(16):320.

100. Leinonen R, Sugawara H, Shumway M. The Sequence Read Archive. Nucleic Acids Res. januari 2011;39(Database issue):D19-21.

101. Choudhary S. pysradb: A Python package to query next-generation sequencing metadata and data from NCBI Sequence Read Archive [Internet]. F1000Research; 2019 [geciteerd 28 februari 2022]. Beschikbaar op: https://f1000research.com/articles/8-532

102. Crusoe MR, Alameldin HF, Awad S, Boucher E, Caldwell A, Cartwright R, e.a. The khmer software package: enabling efficient nucleotide sequence analysis [Internet]. F1000Research; 2015 [geciteerd 28 februari 2022]. Beschikbaar op: https://f1000research.com/articles/4-900

103. Li H. Aligning sequence reads, clone sequences and assembly contigs with BWA-MEM. ArXiv13033997 Q-Bio [Internet]. 26 mei 2013 [geciteerd 28 februari 2022]; Beschikbaar op: http://arxiv.org/abs/1303.3997

104. Dobin A, Davis CA, Schlesinger F, Drenkow J, Zaleski C, Jha S, e.a. STAR: ultrafast universal RNA-seq aligner. Bioinformatics. 1 januari 2013;29(1):15-21.

105. Picard Tools - By Broad Institute [Internet]. [geciteerd 28 februari 2022]. Beschikbaar op: http://broadinstitute.github.io/picard/

106. Li H, Handsaker B, Wysoker A, Fennell T, Ruan J, Homer N, e.a. The Sequence Alignment/Map format and SAMtools. Bioinformatics. 15 augustus 2009;25(16):2078-9.

107. Amemiya HM, Kundaje A, Boyle AP. The ENCODE Blacklist: Identification of Problematic Regions of the Genome. Sci Rep. 27 juni 2019;9(1):9354.

108. Anders S, Pyl PT, Huber W. HTSeq—a Python framework to work with high-throughput sequencing data. Bioinformatics. 15 januari 2015;31(2):166-9.

109. Wang L, Wang S, Li W. RSeQC: quality control of RNA-seq experiments. Bioinformatics. 15 augustus 2012;28(16):2184-5.

110. Zhang Y, Liu T, Meyer CA, Eeckhoute J, Johnson DS, Bernstein BE, e.a. Model-based Analysis of ChIP-Seq (MACS). Genome Biol. 17 september 2008;9(9):R137.

111. Li Q, Brown JB, Huang H, Bickel PJ. Measuring reproducibility of high-throughput experiments. Ann Appl Stat. september 2011;5(3):1752–79.

112. Ritchie ME, Phipson B, Wu D, Hu Y, Law CW, Shi W, e.a. limma powers differential expression analyses for RNA-sequencing and microarray studies. Nucleic Acids Res. 20 april 2015;43(7):e47.

113. Love MI, Huber W, Anders S. Moderated estimation of fold change and dispersion for RNA-seq data with DESeq2. Genome Biol. 5 december 2014;15(12):550.

114. Stephens M. False discovery rates: a new deal. Biostatistics. 1 april 2017;18(2):275–94.

115. Gu Z, Eils R, Schlesner M. Complex heatmaps reveal patterns and correlations in multidimensional genomic data. Bioinforma Oxf Engl. 15 september 2016;32(18):2847–9.

116. Gu Z, Gu L, Eils R, Schlesner M, Brors B. circlize Implements and enhances circular visualization in R. Bioinforma Oxf Engl. oktober 2014;30(19):2811–2.

117. Pagès H, Carlson M, Falcon S, Li N. AnnotationDbi: Manipulation of SQLite-based annotations in Bioconductor [Internet]. Bioconductor version: Release (3.14); 2022 [geciteerd 28 februari 2022]. Beschikbaar op: https://bioconductor.org/packages/AnnotationDbi/

118. Pathview: an R/Bioconductor package for pathway-based data integration and visualization | Bioinformatics | Oxford Academic [Internet]. [geciteerd 28 februari 2022]. Beschikbaar op: https://academic.oup.com/bioinformatics/article/29/14/1830/232698

119. Quinlan AR, Hall IM. BEDTools: a flexible suite of utilities for comparing genomic features. Bioinforma Oxf Engl. 15 maart 2010;26(6):841-2.

120. Chen X, Miragaia RJ, Natarajan KN, Teichmann SA. A rapid and robust method for single cell chromatin accessibility profiling. Nat Commun. 17 december 2018;9(1):5345.

121. Hao Y, Hao S, Andersen-Nissen E, Mauck WM, Zheng S, Butler A, e.a. Integrated analysis of multimodal single-cell data. Cell. 24 juni 2021;184(13):3573-3587.e29.

122. Tirosh I, Izar B, Prakadan SM, Wadsworth MH, Treacy D, Trombetta JJ, e.a. Dissecting the multicellular ecosystem of metastatic melanoma by single-cell RNA-seq. Science. 8 april 2016;352(6282):189-96.

123. Couser NL, redacteur. Preface. In: Ophthalmic Genetic Diseases [Internet]. Philadelphia: Elsevier; 2019 [geciteerd 19 mei 2022]. p. xi. Beschikbaar op: https://www.sciencedirect.com/science/article/pii/B9780323654142050017

124. Khawaja AP, Rojas Lopez KE, Hardcastle AJ, Hammond CJ, Liskova P, Davidson AE, e.a. Genetic Variants Associated With Corneal Biomechanical Properties and Potentially Conferring Susceptibility to Keratoconus in a Genome-Wide Association Study. JAMA Ophthalmol. 1 september 2019;137(9):1005–12.

125. Hardcastle AJ, Liskova P, Bykhovskaya Y, McComish BJ, Davidson AE, Inglehearn CF, e.a. A multi-ethnic genome-wide association study implicates collagen matrix integrity and cell differentiation pathways in keratoconus. Commun Biol. 1 maart 2021;4(1):266.

